# Exploring different virulent proteins of human respiratory syncytial virus for designing a novel epitope-based polyvalent vaccine: Immunoinformatics and molecular dynamics approaches

**DOI:** 10.1101/2022.02.02.478791

**Authors:** Abu Tayab Moin, Md. Asad Ullah, Rajesh B. Patil, Nairita Ahsan Faruqui, Bishajit Sarkar, Yusha Araf, Sowmen Das, Khaza Md. Kapil Uddin, Md Shakhawat Hossain, Md. Faruque Miah, Mohammad Ali Moni, Dil Umme Salma Chowdhury, Saiful Islam

## Abstract

Human Respiratory Syncytial Virus (RSV) is one of the most prominent causes of lower respiratory tract infections (LRTI), contributory to infecting people from all age groups - a majority of which comprises infants and children. The implicated severe RSV infections lead to numerous deaths of multitudes of the overall population, predominantly the children, every year. Consequently, despite several distinctive efforts to develop a vaccine against the RSV as a potential countermeasure, there is no approved or licensed vaccine available yet, to control the RSV infection effectively. Therefore, through the utilization of immunoinformatics tools, a computational approach was taken in this study, to design and construct a multi-epitope polyvalent vaccine against the RSV-A and RSV-B strains of the virus. Potential predictions of the T-cell and B-cell epitopes were followed by extensive tests of antigenicity, allergenicity, toxicity, conservancy, homology to human proteome, transmembrane topology, and cytokine-inducing ability. The most promising epitopes (i.e. 13 CTL epitopes, 9 HTL epitopes, and 10 LBL epitopes) exhibiting full conservancy were then selected for designing the peptide fusion with appropriate linkers, having hBD-3 as the adjuvant. The peptide vaccine was modeled, refined, and validated to further improve the structural attributes. Following this, molecular docking analysis with specific TLRs was carried out which revealed excellent interactions and global binding energies. Additionally, molecular dynamics (MD) simulation was conducted which ensured the stability of the interactions between vaccine and TLR. Furthermore, mechanistic approaches to imitate and predict the potential immune response generated by the administration of vaccines were determined through immune simulations. Owing to an overall evaluation, *in silico* cloning was carried out in efforts to generate recombinant pETite plasmid vectors for subsequent mass production of the vaccine peptide, incorporated within *E.coli*. However, more in vitro and in vivo experiments can further validate its efficacy against RSV infections.

## 1. Introduction

The Human Respiratory Syncytial Virus (hRSV), a member of the family of *Paramyxoviridae,* is known to be the primary cause of lower respiratory tract infections (LRTI), including pneumonia and bronchiolitis, in infants, children, as well as elderly and immunocompromised individuals [1–2]. RSV is an enveloped virus that contains a single-stranded, negative-sense RNA with a genome size of about 15.2 kb. As of yet, two major RSV antigenic subtypes have been identified, RSV-A and RSV-B, exhibiting differential sequence divergence throughout their genome; RSV-A has been seen to be more prevalent than RSV-B [2–3]. Antibody cross-reactivity patterns revealed these two antigenic subgroups (A and B) for RSV, which were then divided into genotypes based on genetic divergence within the highly variable G gene [4–6]. Contributory to the fact that the RSV attachment (G) protein has a central conserved domain (CCD) with a CX3C motif, which has been known to be linked to the generation of protective antibodies, vaccine candidates including the G protein are of considerable interest [7]. A novel genotype of RSV-A, known as RSV-A ON1 was found in Ontario, Canada, in 2010. RSV-A ON1 has a 72-nucleotide duplication at the G Protein’s C terminus [8], which has been linked to an increased risk of pneumonia and lower respiratory tract infections [9]. However, the two subgroups can coexist and thrive, owing to RSV reinfections being common throughout the life of an infected individual, indicating that cross-immunity against distinct strains is only partial [10]. RSV-A infection is commonly followed by RSV-B infection, although the scenario may vary upon several factors [11]. A claim owing to the antigenic diversity of the G protein states that, both within and between antigenic subgroups, this prominent diversity aids in evading pre-existing host immune responses [12, 13].

RSV severely affects immunocompromised infants and the geriatric population with weaned immune systems. The implicated virus infection is considered globally to be the second largest cause of death, in children under one year of age. RSV-associated acute LRTI is responsible for around 33 million serious respiratory infections a year, according to the World Health Organisation (WHO); resulting in more than 3 million hospitalizations and about 60,000 deaths of children under 5 years of age, and 6.7% of all deaths in infants younger than one-year-old. About a half of these hospitalizations and deaths have since been confirmed to be in infants younger than 6 months of age [14]. Additionally, RSV was identified as the third leading cause of fatal childhood pneumonia after *Streptococcus pneumonia* and *Haemophilus influenza* in 2005, responsible for approximately 66,000 to 199,000 deaths from pneumonia in children younger than 5 years [1]. The consequential impact of RSV on older people may be similar to that of influenza, according to epidemiological research, both in the community and in long-term care institutions. In nursing facilities, attack rates are around 5–10 percent per year, with pneumonia (10–20 percent) and mortality (2–5 percent) being quite common. Moreover, RSV infections cause around 10,000 deaths yearly among those aged 64 and over, according to estimates based on US healthcare databases and viral surveillance results [15]. RSV infects the cells lining the human respiration pathway, including the ciliated epithelial cells, and causes upper and lower respiratory tract complications. Influenza-like diseases and LRTI display clinical symptoms of serious RSV infection. However, the most frequent and serious occurrence of infection in younger children is bronchiolitis. Also, over the lifespan of adults, reinfection by the same and separate strains of RSV is considered to be normal, and therefore, RSV is often termed as a chronic virus [2].

Consequently, over the past two decades, RSV has become a major focus for vaccination studies to decrease the morbidity of lower respiratory tract infections. Several vaccinations and antiviral drugs have been formulated and implemented over the years since its identification, and while multiple vaccines, prophylactic and monoclonal antibody candidates are available in clinical trials, no approved RSV vaccine is currently available to counter RSV [16]. The first RSV vaccine, composed of the formalin-inactivated virus (FI-RSV) from the Bernett strain, was studied in a clinical trial back in 1966. Unfortunately, however, the FI-RSV vaccine had a disastrous effect as it struggled to induce an effective neutralizing antibody response, thus preventing infection [2]. Large quantities of eosinophils were discovered in the lungs of children and infants with severe illness, but not in individuals who had a normal RSV infection. Following this unanticipated outcome, it was crucial to design a safe RSV vaccine, which included increased testing for vaccine-induced illnesses [17–21]. The inability of the vaccine to elicit effective neutralizing antibodies or memory CD8+ T cells, as well as the production of a significant inflammatory CD4 T cell response, contributed to this vaccine-induced illness [22–25].

Numerous modified RSV vaccine candidates have been designed after the failure of the FI-RSV vaccine trial and many of them are now in clinical trials. However, none of those vaccine candidates being licensed have so far made it to the international economy, for mass production and administration. Although live-attenuated vaccines can stimulate both a humoral and cellular immune response, clinical trials have revealed some potential drawbacks. Chimpanzees are used to compare the amount of attenuation of vaccinations that are candidates for use in humans. Karron et al. found that RSV vaccines that were temperature sensitive and had a high degree of attenuation in chimps could cause infection in the lower respiratory tract in children [26]. Furthermore, recombinant vector-based vaccinations allow for the presentation of one or more antigens encoded on a viral vector such as PIV3 or adenovirus [27]. Intranasal delivery of a new BLP (bacterial-like particle) conjugated to the RSV fusion protein stimulates both mucosal IgA responses and increased IFN-production in a different sort of vaccination approach [28]. Although both represent effective approaches, further assays to evaluate the long-lasting immune responses are paramount [29].

In addition, it has been observed that, with the use of RSV vaccine candidates, palliative treatment with RSV anti-infective drugs is also required [30]. Merely two approved RSV antivirals are currently available, which include, palivizumab, a humanized preventive monoclonal antibody, and aerosolized ribavirin for therapy. The symptoms of RSV infections can be alleviated by these two antivirals, although they cannot serve prophylactic measures [31]. While studies are underway to identify an effective antiviral therapy or countermeasure to prevent RSV spread and infection, these studies have not been able to deliver any satisfactory findings that can be used to tackle RSV infections [32].

The production of a viable vaccine candidate against a specific pathogen by traditional means can often take many years [33]. However, the age of vaccine production, especially the novel epitope-based “subunit vaccines,” has been enriched by today’s modern technology and the availability of genomic information for almost all pathogens. These subunit vaccines consist only of the antigenic protein segments of the target pathogen and hence, toxic and immunogenic or allergenic parts of the antigen can be dissipated during the construction of the specific vaccine [34]. Again, the development of vaccines using these computer-based approaches takes far less time, and thus greatly reduces the expense of construction and development [35, 36].

The immunoinformatics approach in this study was used to establish successful polyvalent vaccines against the virulent strains of both forms of RSV, i.e. RSV-A and RSV-B respectively. Immunoinformatics is a vaccine modeling process that allows predictions using several computational methods. The novel antigens of a pathogen or virus are identified in immunoinformatics by dissecting its genomic data and then, through the utilization of various *in silico* biology and bioinformatics tools for vaccine design and development, by analyzing the target pathogen genome [35, 37]. In our research, a polyvalent epitope-based vaccine blueprint was produced that could produce a significant immune response to both RSV-A and RSV-B forms, targeting the phosphoprotein (P protein), nucleoprotein (N protein), fusion glycoprotein (F protein), and major surface glycoprotein (mG protein) of these viruses. Since RSV-A is more prevalent than RSV-B, as a model, the vaccine was developed using RSV-A [2]. For the T-cell and B-cell epitope prediction, the RSV-A P protein, N protein, F protein, and mG protein were used and then the epitopes with 100 % conservancy in both species along with some other selection criteria were selected for vaccine construction. The criteria for selecting the epitopes include i.e., antigenicity (the parameter that measures whether the epitopes stimulate a high antigenic response), non-allergenicity (to ensure that the epitopes do not cause any unintended allergic reaction inside the body), non-toxicity, conservancy across the selected organisms, as well as non-homologation of the human proteome. It is, therefore, expected that the vaccine will be effective against both the subtypes - RSV-A and RSV-B. The most common vaccine target for RSV is known to be the F protein [574 amino acids (aa) in length], which is a highly conserved protein in both RSV forms. The F protein mediates the fusion and attachment of the virus to its target cells along with the mG protein, thus facilitating viral entry [2, 38]. The F1 (aa 137–574) and F2 (aa 1–109) subunits form a homotrimer in the mature F protein, and the F1 subunit is required for membrane fusion. The F protein has two different conformations i.e., the pre-fusion and post-fusion conformations [39, 40]. The protein rearranges to a more stable post-fusion form during infection to allow viral entrance into the host cell. Antibodies having neutralizing activity identify at least two antigenic sites on both the pre-fusion and post-fusion forms of F (sites II and IV) [41–43]. In this study, the precursor F0 protein was targeted to retrieve all of the potential antigenic epitopes. The possible conformational change of the F protein, as well as the cleavage sites of the protein sequence, were taken into account while generating the potential epitopes [39, 40].

The viral genome of RSV is surrounded by N protein, and the P protein is a vital component of the viral RNA-dependent RNA polymerase complex which is necessary for the proper replication and transcription of RSV [44]. Therefore, in our study, these four proteins were used as possible targets to design a vaccine to suppress these viral proteins, preventing viral entry, and thus interfering with the life cycle of the virus.

## 2. Methods and Materials

The high throughput immunoinformatics and MD approaches of vaccine designing are illustrated in a step-by-step processes in **Fig 1**.

**Fig 1.**
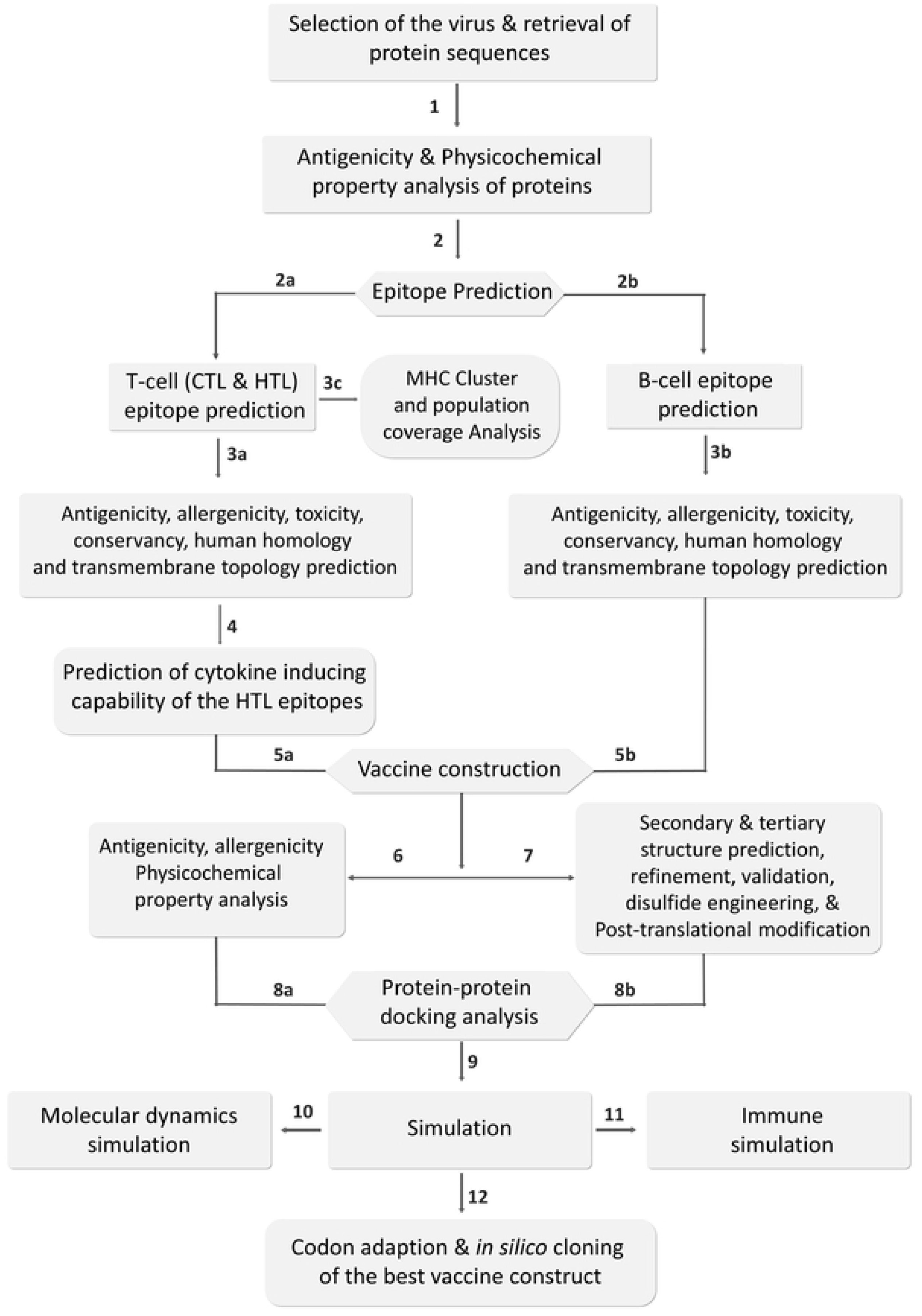
The step-by-step procedures of immunoinformatics and molecular dynamics approaches used in the vaccine designing study.

### 2.1. Protein sequences identification and retrieval

Through existing literature reviews in the National Center for Biotechnology Information (NCBI) (https://www.ncbi.nlm.nih.gov/) database, the RSV-A and RSV-B viruses were identified and selected along with their target proteins (i.e., P protein, N protein, F protein, and mG protein). The sequences of target proteins of the selected strains (i.e., RSV strain A2 and RSV strain B1) were then extracted from the UniProt (https://www.uniprot.org/) database in FASTA format. The NCBI Protein database is a collection of SwissProt, PIR, PRF, and PDB sequences. It also includes GenBank, RefSeq, and TPA translations from elucidated coding regions.

### 2.2. Prediction of antigenicity and analysis of physicochemical properties of the selected proteins

Using the online antigenicity prediction tool, VaxiJen v2.0 (http://www.ddg-pharmfac.net/vaxijen/VaxiJen/VaxiJen.html), the antigenicity of the target protein sequences was predicted with the prediction precision parameter threshold kept at 0.4. This tool uses the method of transformation of auto cross-covariance (ACC) to predict the antigenicity of query proteins or peptides and provides results with an accuracy of 70% to 89%. For this reason, this server is the widely used and accepted server to determine the antigenicity of query proteins [45]. ProtParam tool of the ExPASy server (https://web.expasy.org/protparam/) has subsequently determined numerous physicochemical properties, i.e. the number of amino acids, molecular weight, number of total atoms, theoretical pI, instability index, extinction coefficient, half-life, grand average of hydropathicity (GRAVY), etc. of the target proteins [46].

### 2.3. Prediction of T-cell and B-cell epitopes

The two major types of T-cells, cytotoxic T-cells, and Helper T-cells are both considered essential for the successful design of the vaccine [47]. For specific antigen recognition of the major histocompatibility complex class I (MHC-I) or CD8+ cytotoxic T-lymphocytic (CTL) epitopes on the surface of the antigen-presenting cells (APCs), the cytotoxic T-cells are important. Additionally, the helper T-cells are considered to be a crucial component of adaptive immunity that interacts on the surface of APCs with major histocompatibility complex class II (MHC-II) or CD4+ helper T-lymphocytic (HTL) epitopes. They function in activating the B-cell, macrophages, and even cytotoxic T-cells [48, 49]. On the other hand, B-cells produce antigen-specific immunoglobulins after their activation [50]. They can identify solvent-exposed antigens via membrane-bound immunoglobulins called B cell receptors (BCRs) [51]. B-cell epitopes are important for defense against viral infections because they are the essential immune system components that activate an adaptive immune response in response to a specific viral infection. Therefore, the B-cell epitopes are used as one of the crucial building blocks of the subunit vaccine. There are two types of B-cell epitopes: linear B-cell epitopes (LBL) and conformational B-cell epitopes, also known as continuous and discontinuous B-cell epitopes, respectively [52].

The T-cell and B-cell epitope prediction was performed using the Immune Epitope Database or IEDB (https://www.iedb.org/), which contains extensive experimental data on antibodies and epitopes [53]. For the prediction of MHC Class-I or CTL epitopes for several human leukocyte antigen (HLA) alleles, i.e., HLA A*01:01, HLA A*03:01, HLA A*11-01, HLA A*02:01, HLA A*02:06, and HLA A*29:02, the recommended IEDB NetMHCpan 4.0 prediction method was used. The default prediction method selection of the server is ‘IEDB recommended’ which utilizes the best available technique for a specific MHC molecule based on the availability of predictors and observes the predicted performance for a specific allele. It is updated regularly based on predictor availability. NetMHCpan EL 4.1 is currently used across all alleles for peptide: MHC Class-I binding prediction. Again, for the prediction of MHC class-II or HTL epitopes for DRB1*03:01, DRB1*04:01, DRB1*15:01, DRB3*01:01, DRB5*01:01, and DRB4*01:01 alleles, the recommended IEDB 2.22 prediction method was used. If any corresponding predictor is available for the MHC molecule, the IEDB recommended method employs the Consensus method, combining NN-align, SMM-align, CombLib, and Sturniolo; otherwise, NetMHCIIpan is used. If any three of the four approaches are available, the Consensus approach evaluates them all, with Sturniolo as the final option. Henceforth, based on their ranking, the top-scored HTL and CTL epitopes that were found to be common for all of the selected corresponding HLA alleles were considered for further analyses. All the parameters were retained by opting for default during the T-cell epitope prediction. Subsequently, B-cell epitopes of the proteins were predicted using the BepiPred linear epitope prediction method 2.0, maintaining all the default parameters. Using a Random Forest algorithm trained on epitope and non-epitope amino acids obtained from crystal structures, the BepiPred-2.0 server predicted linear B-cell epitopes from a protein sequence. Following this, a sequential prediction smoothing was conducted. Residues with scores greater than the threshold (default value of 0.5) were thought to constitute epitopes [54]. Finally, the top-scored LBL epitopes containing more than ten amino acids were primarily regarded as potential candidates for further analysis.

Conformational or discontinuous B-cell epitopes are critical components to induce antibody-mediated humoral immunity within the body. While designing a vaccine, efficient conformational B-cell epitopes should be included alongside the LBLs to elicit a better immunogenic response against the pathogen. The conformational B-cell epitopes of the modeled 3D structure of the vaccine were predicted using IEDB ElliPro, an online server (http://tools.iedb.org/ellipro/) using the default parameters of a minimum score of 0.5 and a maximum distance of 6 angstroms [55]. ElliPro uses three algorithms to predict the protein shape as an ellipsoid, measure the residue PI, and estimate adjacent cluster residues based on their protrusion index (PI) values [56]. ElliPro calculates a score for each output epitope based on an average PI value over the residues of each epitope. Protein residues are contained in 90% of ellipsoids with a PI value of 0.9, while 10% of residues are outside ellipsoids. The center of residue mass residing outside the largest ellipsoid possible was used to calculate the PI value for each epitope residue [57].

### 2.4. Assessment of antigenicity, allergenicity, toxicity, and topology prediction of the epitopes

In this step, several methods for predicting their conservancy, antigenicity, allergenicity, and toxicity were used to evaluate the initially predicted T-cell and B-cell epitopes. To assess the conservancy of the chosen epitopes [58], the conservancy prediction method of the IEDB server (https://www.iedb.org/conservancy/) was used. Additionally, the components of the vaccine should be highly antigenic, non-allergenic at the same time, and also devoid of toxic reactions. In this step, the antigenicity determination tool VaxiJen v2.0 (http://www.ddg-pharmfac.net/vaxijen/VaxiJen/VaxiJen.html) was used again for the determination of antigenicity [45]. Two different tools were then used, i.e. AllerTOP v2.0 (https://www.ddg-pharmfac.net/AllerTOP/) and AllergenFP v1.0 (http://ddg-pharmfac.net/AllergenFP/) to obtain the highest precision for prediction of allergenicity. Both of the tools are based on auto cross-covariance (ACC) transformation of protein sequences into uniform equal-length vectors. However, the AllerTOP v2.0 server has a better 88.7 % prediction accuracy than the AllergenFP v1.0 server (87.9 %) [59, 60]. In addition, the ToxinPred (http://crdd.osdd.net/raghava/toxinpred/) server was used to predict toxicity for all epitopes by using the Support Vector Machine (SVM) prediction method to keep all the default parameters. The SVM is a widely accepted machine learning technique for toxicity prediction since it can differentiate the toxic and non-toxic epitopes quite efficiently [61]. Finally, using the TMHMM v2.0 server (http://www.cbs.dtu.dk/services/TMHMM/), the transmembrane topology prediction of all the epitopes was performed to predict whether the epitopes were exposed inside or outside, keeping the parameters at their default values. TMHMM uses an algorithm called N-best (or 1-best in this case) to predict the most probable location and orientation of transmembrane helices in the sequence [62].

### 2.5. Cytokine inducing capacity prediction of the epitopes

Several cytokine types, including IFN-γ, IL-4 (interleukin-4), and IL-10 (interleukin-10) are produced by helper T cells to activate various immune cells, i.e. cytotoxic T cells, macrophages, etc. [63]. As a result, it is crucial to know whether HTL epitopes are capable of producing key cytokines to induce an immune response against the virus before designing a vaccine. The induction capacity of the predicted HTL epitopes for interferon-γ (IFN-γ) was determined using the IFNepitope (http://crdd.osdd.net/raghava/ifnepitope/) server. Based on analyzing a dataset that includes IFN-γ inducing and non-inducing peptides, the server determines the probable IFN-γ inducing epitopes. To determine the IFN-γ inducing capacity, the Design module and the Hybrid (Motif + SVM) prediction approach were used. The Hybrid prediction approach is considered to be a highly precise approach to the prediction of the epitope-inducing capacity of IFN-γ [64]. In addition, IL-4 and IL-10 inducing HTL epitope properties were determined using the servers IL4pred (https://webs.iiitd.edu.in/raghava/il4pred/index.php) and IL10pred (http://crdd.osdd.net/raghava/IL-10pred/) [65, 66]. The SVM method was used on both servers, where the default threshold values were kept at 0.2 and -0.3, respectively.

### 2.6. Conservancy and human proteome homology prediction

The conservancy analysis of the specified epitopes was conducted using the IEDB server’s epitope conservancy analysis module (https://www.iedb.org/conservancy/) [58]. The epitopes that were found to be fully conserved among the selected strains were taken for the construction of the vaccine since this will ensure and facilitate the broad-spectrum activity of the polyvalent vaccine over the two selected RSV species or types. The homology of the human proteome epitopes was determined by the BLAST (BlastP) protein module of the BLAST tool (https://blast.ncbi.nlm.nih.gov/Blast.cgi), where Homo sapiens (taxid:9606) was used for comparison, keeping all other default parameters. An e-value cut-off of 0.05 was set and epitopes were selected as non-homologous pathogen peptides that showed no hits below the e-value inclusion threshold [67]. The epitopes found to be highly antigenic, non-allergenic, non-toxic, fully conserved, and non-homologous to the human proteome were considered among all the initially selected epitopes to be the best-selected epitopes or the most promising epitopes, and only these selected epitopes were used in the construction of the vaccine.

### 2.7. Population coverage and cluster analyses of the epitopes and their MHC alleles

A crucial requirement is to consider the distribution of unique HLA alleles among the different populations and ethnicities around the world to design a multi-epitope vaccine since the expression of different HLA alleles can vary from population to population. The IEDB resource for population coverage (http://tools.iedb.org/population/) was used for analyzing the population coverage of the most promising epitopes across several HLA alleles in various regions around the world. Denominated MHC restriction of T cell responses and polymorphic HLA combinations were considered in the analysis. All the parameters were maintained at their default conditions during the study.

Furthermore, the human MHC genomic region or HLA is enormously polymorphic, with thousands of alleles; many of which code for a different molecule. MHCcluster is a program that organizes the MHC molecules into functional clusters based on their predicted binding specificity. The approach provides a user-friendly online interface that allows the user to include any MHC in the analysis. A static heat map and graphical tree-based visualizations of the functional relationship between the MHC variants are included in the output as well as a dynamic TreeViewer interface that displays both the functional relationship and the individual binding specificities of the MHC molecules [68]. To evaluate the relationship between the selected MHC alleles, cluster analysis of the MHC alleles was done using the online tool MHCcluster 2.0 https://services.healthtech.dtu.dk/service.php?MHCcluster-2.0). During the study, 50,000 peptides to be used were retained, 100 bootstrap measurements were retained, and both HLA super-type (MHC Class-I) and HLA-DR (MHC class-II) members were chosen.

### 2.8. Designing of the multi-epitope subunit vaccine

The most promising antigenic epitopes have been linked with each other to create a fusion peptide using an adjuvant and linkers. Human beta-defensin-3 (hBD-3) used an adjuvant sequence that was linked to the epitopes by EAAAK linkers. Adjuvants are considered to play important roles in improving the antigenicity, immunogenicity, stability, and durability of the developed vaccine. The hBD-3 plays a vital role in host immune responses against the pathogens (i.e., innate mucosal defense within the respiratory tract) and is highly significant against respiratory infections [69–71].

The epitopes were also appended to the pan HLA-DR epitope (PADRE) sequence. By enhancing the ability of CTL vaccine epitopes, the PADRE sequence activates the immune responses [34]. In the conjugation of the CTL, HTL, and LBL epitopes, the AAY, GPGPG, and KK linkers were used, respectively. The EAAAK linkers have a viable partition of bifunctional fusion protein domains [72], while the GPGPG linkers are ideal for preventing junctional epitope production and optimizing the processing and presentation of the immune system [73]. The AAY linker is also commonly used in the design trials of the *in silico* vaccine since this linker offers successful and efficient epitope conjugation [74]. In addition, bi-lysine (KK) linkers are active in the autonomous immunological function of vaccine epitopes [75].

### 2.9. Physicochemical property analyses of the vaccine with antigenicity and allergenicity test

To build a timely and successful immune response to the pathogenic attack, the constructed vaccine should be strongly antigenic. The antigenicity of the vaccine model was estimated using VaxiJen v2.0 (http://www.ddg-pharmfac.net/vaxijen/VaxiJen/VaxiJen.htm), keeping the threshold value fixed at 0.4 [45]. The findings of the Vaxijen v2.0 server were further cross-checked by the ANTIGENpro module of the SCRATCH protein predictor (http://scratch.proteomics.ics.uci.edu/), holding all the default parameters [76], to achieve better prediction precision. Three separate online methods have estimated the allergenicity of the vaccine structures, i.e. AlgPred (http://crdd.osdd.net/raghava/algpred/), AllerTop v2.0 (https://www.ddgpharmfac.net/AllerTOP/) and AllergenFP v1.0 (http://dg-pharmfac.net/AllergenFP/), to ensure optimum prediction precision. The AlgPred (http://crdd.osdd.net/raghava/algpred/) server aims to combine multiple allergenicity determination methods to reliably determine possible allergenic proteins [77, 78]. To predict the vaccine’s allergenicity, the MEME/MAST motif prediction approach was used. The physicochemical properties of the built vaccine were then estimated by the same online instrument, ProtParam (https://web.expasy.org/protparam/)[15], which was previously utilized. The solubility of vaccine constructs was also estimated alongside the physicochemical property study by the SOLpro module of the SCRATCH protein predictor (http://scratch.proteomics.ics.uci.edu/) and later further explained by the Protein-Sol server (https://protein-sol.manchester.ac.uk/). The solubility of a query protein is predicted by all these servers with remarkable precision. SolPro produces its predictions based on the SVM method, while Protein-Sol uses a rapid method of deciding the results based on sequence [76, 79]. All the parameters of the servers were maintained at their default values during the solubility review.

### 2.10. Secondary and tertiary structure prediction of the vaccine construct

The vaccine construct was subjected to secondary structure prediction following physicochemical analysis. For this, several online resources were used to preserve all the default parameters, i.e. PSIPRED (http://bioinf.cs.ucl.uk/psipred/) (using the PSIPRED 4.0 prediction method), GOR IV (https://npsa-prabi.ibcp.fr/cgibin/npsaautomat.pl?page=/NPSA/npsagor4.html), SOPMA (https://npsa-prabi.ibcp.fr/cgibin/npsaautomat.pl?page=/NPSA/npsasopma.html) and SIMPA96 (https://npsaprabi.ibcp.fr/cgibin/npsaautomat.pl?page=/NPSA/npsanpsa.html) and SIMPA96 (https://npsa.npsa.npsa.npsa.npsa. To predict the percentages or quantities of amino acids in α helix, β-sheet, and coil structure formations, these servers are considered to be reliable, quick, and effective [80–84]. Moreover, determination of the tertiary or 3D structure of the vaccine construct was carried out using the RaptorX online server (http://raptorx.uchicago.edu/). Using an easy and powerful template-based method [85], the server predicts the tertiary or 3D structure of a query protein. Furthermore, RaptorX uses a deep learning method to enable distance-based protein folding. This server has also been rated first in contact prediction in both CASP12 and CASP13, making it an ideal server for 3D structure determination [86].

### 2.11. Refinement and validation of tertiary structure of the vaccine

The tertiary structure prediction of the proteins using computational methods also requires extensive refinement, to turn predicted models with lower resolution into models that closely match the native protein structure. Therefore, a GalaxyWEB server (http://galaxy.seoklab.org/) using the GalaxyRefine module further refined the created tertiary structure of the proposed vaccine model. The server uses dynamic simulation and the refinement approach is tested by CASP10 to refine the tertiary protein structures [87, 88]. Furthermore, validation of the refined protein was carried out by analyzing the Ramachandran plot created by the PROCHECK (https://servicesn.mbi.ucla.edu/PROCHECK/) tool [89, 90]. Along with PROCHECK for protein validation, another online platform, ProSA-web (https://prosa.services.came.sbg.ac.at/prosa.php) was also used. A z-score that expresses the consistency of a query protein structure is created by the PROCHECK server. In the latest PDB database, a z-score residing within the z-score spectrum of all experimentally defined protein chains represents a higher consistency of the query protein [91].

### 2.12. Vaccine protein disulfide engineering analysis

Disulfide bonds are more likely to form in a few regions within a protein structure, providing stability through reduced conformational entropy and increased free energy concerning the denatured state. However, disulfide engineering is the process of introducing disulfide bonds to a target protein to increase its stability. In this experiment, the Disulfide by Design (DbD)2 v12.2 (http://cptweb.cpt.wayne.edu/DbD2/) online tool was used to predict the locations and further design the disulfide bonds within the vaccine proteins [92]. The tool was developed using computational approaches to predict the protein structure [93, 94], and the algorithm of this server accurately estimates the χ3 torsion angle based on 5the Cβ–Cβ distance using a geometric model derived from native disulfide bonds. The Caf-Cβ-Sγ angle is allowed some tolerance in the DbD2 server based on the wide range found in native disulfides. To facilitate the ranking process, DbD2 estimates an energy value for each potential disulfide and mutant PDB files may be generated for selected disulfides [95].

The χ3 angle was held at -87 ° or +97 ° ±10 during the experiment to cast off various putative disulfides that were generated using the default angles of +97 ° ±30 ° and -87 ° ±30 °. Additionally, the angle of Caf-Cβ-Sγ was set to its default value of 114.6° ±10. Finally, to allow disulfide bridge formation, residue pairs with energy less than 2.2 Kcal/mol were selected and mutated to cysteine residue [96]. The energy value of 2.2 Kcal/mol was chosen as the disulfide bond selection threshold since 90% of native disulfide bonds are usually considered to have an energy value of less than 2.2 Kcal/mol [92].

### 2.13. Post-translational modification analysis

For posttranslational modification analysis of the vaccine construct comprising of the B-cell and T-cell epitopes, the NetNGlyc-1.0 (http://www.cbs.dtu.dk/services/NetNGlyc-1.0), NetOGlyc4.0 (http://www.cbs.dtu.dk/services/NetOGlyc-4.0), and NetPhos-3.1 (http://www.cbs.dtu.dk/services/NetPhos-3.1) servers were utilized. The NetNglyc server uses artificial neural networks to predict N-glycosylation sites in human proteins by examining the sequence context of Asn-Xaa-Ser/Thr sequons [97]. Any potential that exceeds the default threshold of 0.5 indicates a predicted glycosylated site. The average output of nine neural networks is used to get the ’potential’ score. The NetOglyc server (http://www.cbs.dtu.dk/services/NetOGlyc-4.0) predicts mucin type GalNAc O-glycosylation sites in mammalian proteins using neural networks [98].

This server provides a list of probable glycosylation sites for each input sequence, together with their positions in the sequence and prediction confidence scores. Only locations with a score greater than 0.5 are expected to be glycosylated and the string “POSITIVE” is added to the remark box. Using ensembles of neural networks, the NetPhos 3.1 server (http://www.cbs.dtu.dk/services/NetPhos-3.1) predicts serine, threonine, or tyrosine phosphorylation sites in eukaryotic proteins. Predictions are made for both generic and kinase-specific kinases. A prediction score greater than 0.5 indicates a positive prediction.

### 2.14. Analysis of protein-protein docking

The vaccine protein was docked against several toll-like receptors (TLRs) in protein-protein docking analysis. A strong binding affinity should be present between the vaccine and the TLRs. This is crucial because, after identifying the vaccine that resembles the initial viral infections, TLR proteins generate possible immune responses, and thus help to produce immunity against the pathogen [99]. In this study, different TLRs have been docked with the vaccine protein, i.e. TLR-1 (PDB ID: 6NIH), TLR-2 (PDB ID: 3A7C), TLR-3 (PDB ID: 2A0Z), TLR-4 (PDB ID: 4G8A), and TLR9 (PDB ID: 3WPF). ClusPro v2.0 (https://cluspro.bu.edu/login.php) was used to conduct the docking, where the lower energy score corresponds to the stronger binding affinity. Based on the following equation, the ClusPro server calculates the energy score:

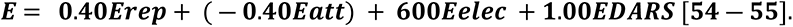

The repulsions and attraction energies owing to van der Waals interactions are denoted by E_rep_ and E_attr_, respectively, whereas E_elec_ signifies the electrostatic energy component. The Decoys’ pairwise structure-based potential is represented by E_DARS_ as the Reference State (DARS) method. Furthermore, another round of docking was carried out using the ZDOCK server which is a rigid-body protein-protein docking tool that employs a combination of shape complementarity, electrostatics, and statistical potential terms for scoring and uses the Fast Fourier Transform algorithm to enable an efficient global docking search on a 3D grid. In the most current benchmark version (Accelerating protein docking in ZDOCK utilizing an advanced 3D convolution library), ZDOCK achieves high predictive accuracy on protein-protein docking benchmarks, with >70 % success in the top 1000 predictions for rigid-body instances [100].

### 2.15. Molecular dynamics simulation studies and MM-PBSA calculations

The docked complexes from the ZDOCK server were used in MD simulations. The complexes being protein-protein in nature with multiple chains, the MD simulations were computationally expensive and performed on the HPC cluster at Bioinformatics Resources and Applications Facility (BRAF), C-DAC, Pune with Gromacs 2020.4 [101] MD simulation package. The CHARMM-36 force field parameters [102, 103] were employed to prepare the topology of protein chains. The system of each TLR along with the bound vaccine was solvated with the single point charge water model [104] in the dodecahedron unit cells and neutralized with the addition of Na^+^ or Cl^-^ counter-ions. The solvated systems were initially energy minimized to relieve the steric clashes if any with the steepest descent criteria until the threshold (Fmax< 10 kJ/mol) was reached. These energy minimized systems were then equilibrated at constant volume and temperature conditions 300 K using modified Berendsen thermostat [105] and then at constant volume and pressure Berendsen barostat [106] for 100 ps each. The equilibrated systems were later subjected to 100 ns production phase MD simulations, where the modified Berendsen thermostat and Parrinello-Rahman barostat [107] were used with covalent bonds restrained using the LINCS algorithm [108]. The long-range electrostatic interaction energies were measured with the cut-off of 12 Å, with the Particle Mesh Ewald method (PME) [109]. The resulting trajectories were analyzed for root mean square deviations (RMSD) in protein backbone atoms, root mean square fluctuations (RMSF) in the side chain atoms of individual chains in each protein complex, the radius of gyration (Rg), and several hydrogen bonds formed between vaccine protein chain and the respective TLR protein chain.

### 2.16. Immune simulation studies

To forecast the immunogenicity and immune response profile of the proposed vaccine, an immune simulation analysis was performed. For the immune simulation study, the C-ImmSim server (http://150.146.2.1/CIMMSIM/index.php) was used to predict real-life immune interactions using machine learning techniques and PSSM (Position-Specific Scoring Matrix) [110]. During the experiment, all the variables except for the time steps were kept at their default parameters. However, the time steps at 1, 84, and 170 were retained (time step 1 is injection at time = 0), and the number of simulation steps was set to 1050. Thus, three injections at four-week intervals were administered to induce recurrent antigen exposure [111].

### 2.17. Codon adaptation and *in silico* cloning within *E.coli* System

Codon adaptation and *in silico* cloning are two significant steps that are conducted to express multi-epitope vaccine construction within an *Escherichia coli* (*E.coli)* K12 strain. In different organisms, an amino acid can be encoded by more than one codon, which is known as codon bias wherefore, the codon adaptation study is carried out to predict an appropriate codon that effectively encodes a specific amino acid in a specific organism. Java Codon Adaptation Tool or JCat server (http://www.jcat.de/) was used for codon optimization [112], and the optimized codon sequence was further analyzed for expression parameters, codon adaptation index (CAI), and GC-content %. The optimum CAI value is 1.0, while a score of > 0.8 is considered acceptable, and the optimum GC content ranges from 30 to 70% [113]. For *in silico* cloning simulation, the pETite vector plasmid was selected which contains a small ubiquitin-like modifier (SUMO) tag as well as a 6x polyhistidine (6X-His) tag, which will facilitate the solubilization and affinity purification of the recombinant vaccine construct [114]. Also, 6X-His can facilitate the swift detection of the recombinant vaccine construct in immunochromatographic assays [115]. The vaccine protein sequence was reverse-translated to the optimized DNA sequence by the server to which EaeI and StyI restriction sites were incorporated at the N-terminal and C-terminal sites, respectively. The newly adapted DNA sequence was then inserted between the EaeI and StyI restriction sites of the pETite vector using the SnapGene restriction cloning software (https://www.snapgene.com/free-trial/) to confirm the expression of the vaccine [116, 117].

### 2.18. Prediction of the vaccine mRNA secondary structure

Two servers, i.e. Mfold (http://unafold.rna.albany.edu/?q=mfold) and RNAfold (http://rna.tbi.univie.ac.at/cgibin/RNAWebSuite/RNAfold.cgi), were used for the mRNA secondary structure prediction. Both of these servers thermodynamically predict the mRNA secondary structures and provide each of the generated structures with minimum free energy (’G Kcal/mol’). The more stable the folded mRNA is, the lower the minimum free energy and vice versa [55][118–120]. To analyze the mRNA folding and secondary vaccine structure, the optimized DNA sequence was first taken from the JCat server and converted via the DNA<->RNA->Protein tool (http://biomodel.uah.es/en/lab/cybertory/analysis/trans.htm) to a possible RNA sequence. The RNA sequence was then gathered from the tool and utilized for prediction into the Mfold and RNAfold servers using the default settings for all the parameters.

## 3. Results

### 3.1. Protein sequences identification and retrieval

From the NCBI database, the RSV viral strains and the query proteins were identified. Following that, the four RSV-A and RSV-B Query Proteins including P protein, N protein, F protein, and mG, were retrieved from the UniProt online database. The UniProt Accession Number and the length of the query proteins are listed in Table 1.

**Table 01.** List of the proteins with their accession numbers used in the vaccine designing study.

| Name of the Virus | Name of the Protein | UniProt Accession Number of the Protein | Length (aa) of the Protein Sequence |
| --- | --- | --- | --- |
| RSV-A | P protein | P03421 | 241 |
|  | N protein | P03418 | 391 |
|  | F protein | P03420 | 574 |
|  | mG protein | P03423 | 298 |
| RSV-B | P protein | O42062 | 241 |
|  | N protein | O42053 | 391 |
|  | F protein | O36634 | 574 |
|  | mG protein | O36633 | 299 |

### 3.2. Prediction of antigenicity and analysis of physicochemical properties of the selected proteins

The selected proteins were analyzed for antigenicity and physicochemical properties through the VaxiJen v2.0 server and ProtParam tool of the ExPASy server, respectively. To be a vaccine candidate, antigenicity is a prerequisite for a protein or amino acid sequence. All of the selected proteins were found to be antigenic in VaxiJen v2.0 server at threshold 0.4. In addition, while P protein and N protein of RSV-A and RSV-B were found to have an acidic theoretical pI (pH lower than 7), F protein and mG protein were found to have a basic theoretical pI (pH higher than 7). The protein having an acidic theoretical pI belongs to negatively charged proteins. Again, in the mammalian cell culture system, all the query proteins had a similar half-life of 30 h and a high aliphatic index (over 60.00) as well. All of the proteins had quite low GRAVY values (lower than -0.909). P protein of the RSV-A and RSV-B had the highest GRAVY value of -0.909 and -0.827, respectively. Whereas, the F protein of RSV-A and RSV-B had the lowest GRAVY value of -0.028 and -0.033, respectively. Furthermore, the F protein of the RSV-A and RSV-B had the highest aliphatic index of 99.97 and 102.35, respectively. **S1 Table** lists the results of the analysis of physicochemical properties of all the query proteins.

### 3.3. Epitope prediction and sorting the most promising epitopes

The RSV-A proteins were selected as models during the prediction of the T-cell and B-cell epitopes by the IEDB server for the construction of the polyvalent vaccine, meaning that the epitopes were selected using only the RSV-A proteins and then only the fully conserved epitopes were taken therefore, the epitopes should confer immunity to the selected strains of both RSV-A and RSV-B. These epitopes were anticipated to induce potential T-cell and B-cell immune responses after the vaccine administration. Based on the ranking, the top CTL and HTL epitopes as well as top B-cell epitopes with lengths over ten amino acids were taken into consideration for further analysis. Following this, a few criteria were selected to filter the best epitopes which included, high antigenicity, non-allergenicity, non-toxicity, conservancy, and human proteome non-homology. Furthermore, the cytokine (i.e., IFN-γ, IL-4, and IL-10) inducing ability of HTL epitopes was also considered to determine whether they can produce at least one of these cytokines. Finally, the epitopes that met these criteria were listed as the most promising epitopes in Table 02 and were later used for the construction of the vaccine. The analysis of transmembrane topology by the TMHMM v2.0 server of the most promising epitopes revealed that PEFHGEDANNR, SFKEDPTPSDNPFS, EVAPEYRHDSPD, VFPSDEFDASISQVNEK, IPNKKPGKKTTTKPTKKPTLKTTKKDPKPQTTKSKEVPTTKP were exposed outside.

**Table 02.**
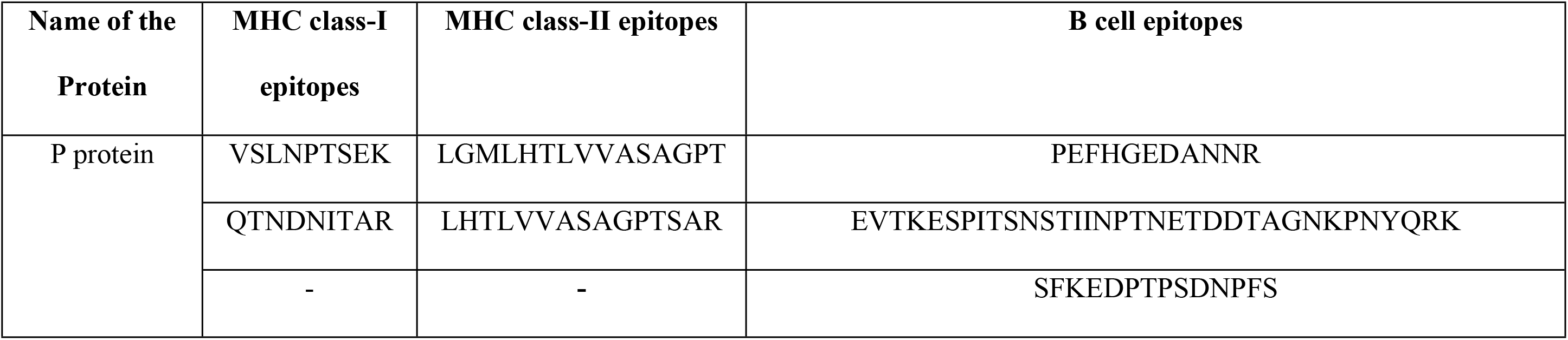
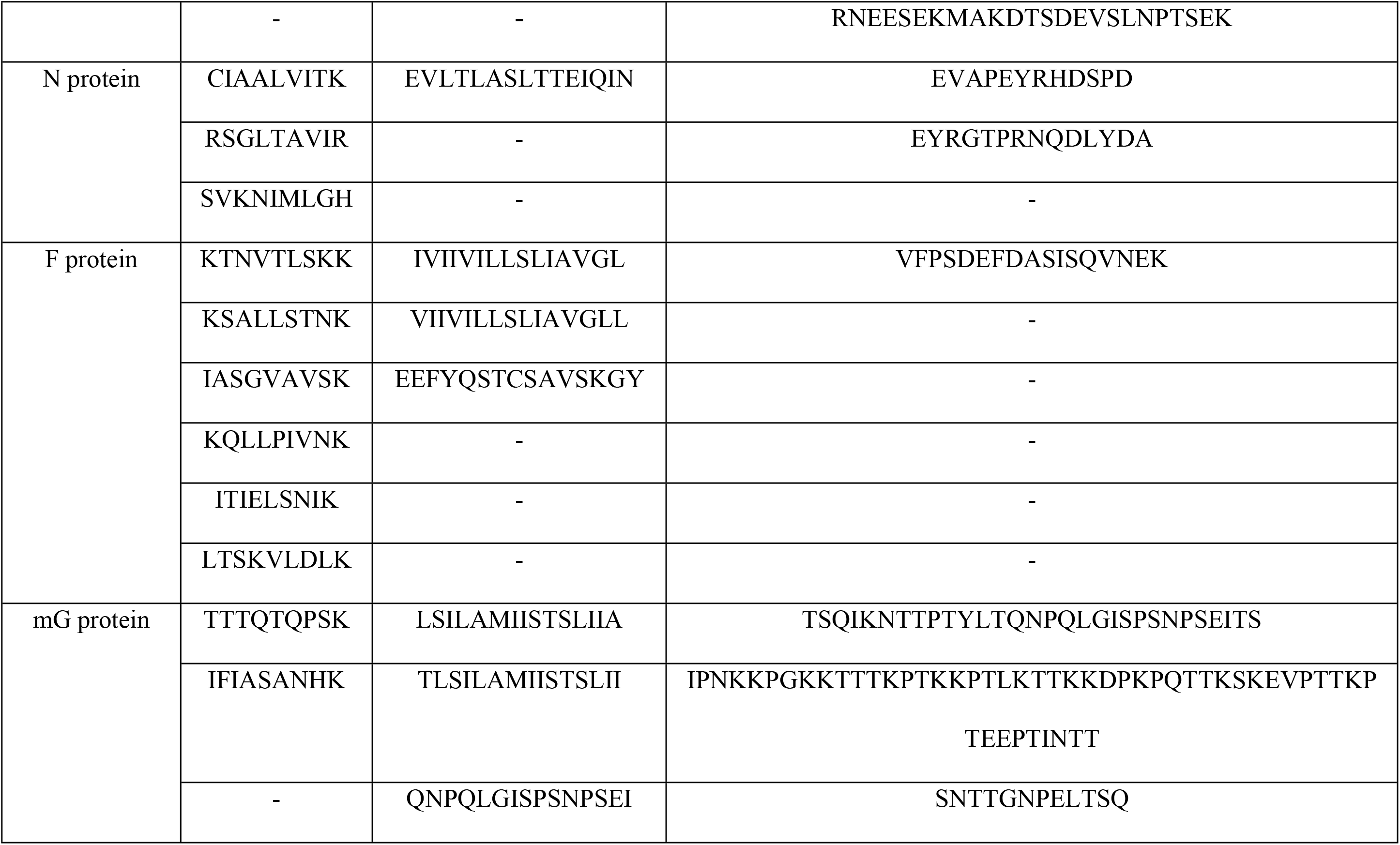
List of the epitopes eventually selected for the construction of the vaccine (selection criteria: antigenicity, non-allergenicity, non-toxicity, 100 % conservancy and non-homolog to the human proteome).

**S2 Table** listed the potential epitopes of P protein and **S3 Table** listed the potential epitopes of N protein. The potential epitopes of F protein are listed in **S4 Table** and the potential epitopes of mG protein are listed in **S5 Table.**

### 3.4. Population coverage and cluster analyses of the epitopes and their MHC alleles

The population coverage analysis showed that 85.70% and 87.92% of the world population were covered by the MHC class-I and class-II alleles and their epitopes, respectively, and 84.62% of the world population was covered by the combined MHC class-I and class-II. While India had the highest percentage of population coverage for the CTL epitopes (87.56 %) as well as HTL epitopes (93.51 %), China had the highest percentage of population coverage for CTL and HTL epitopes in combination (91.80 %) (**Fig 2**).

**Fig 2.**
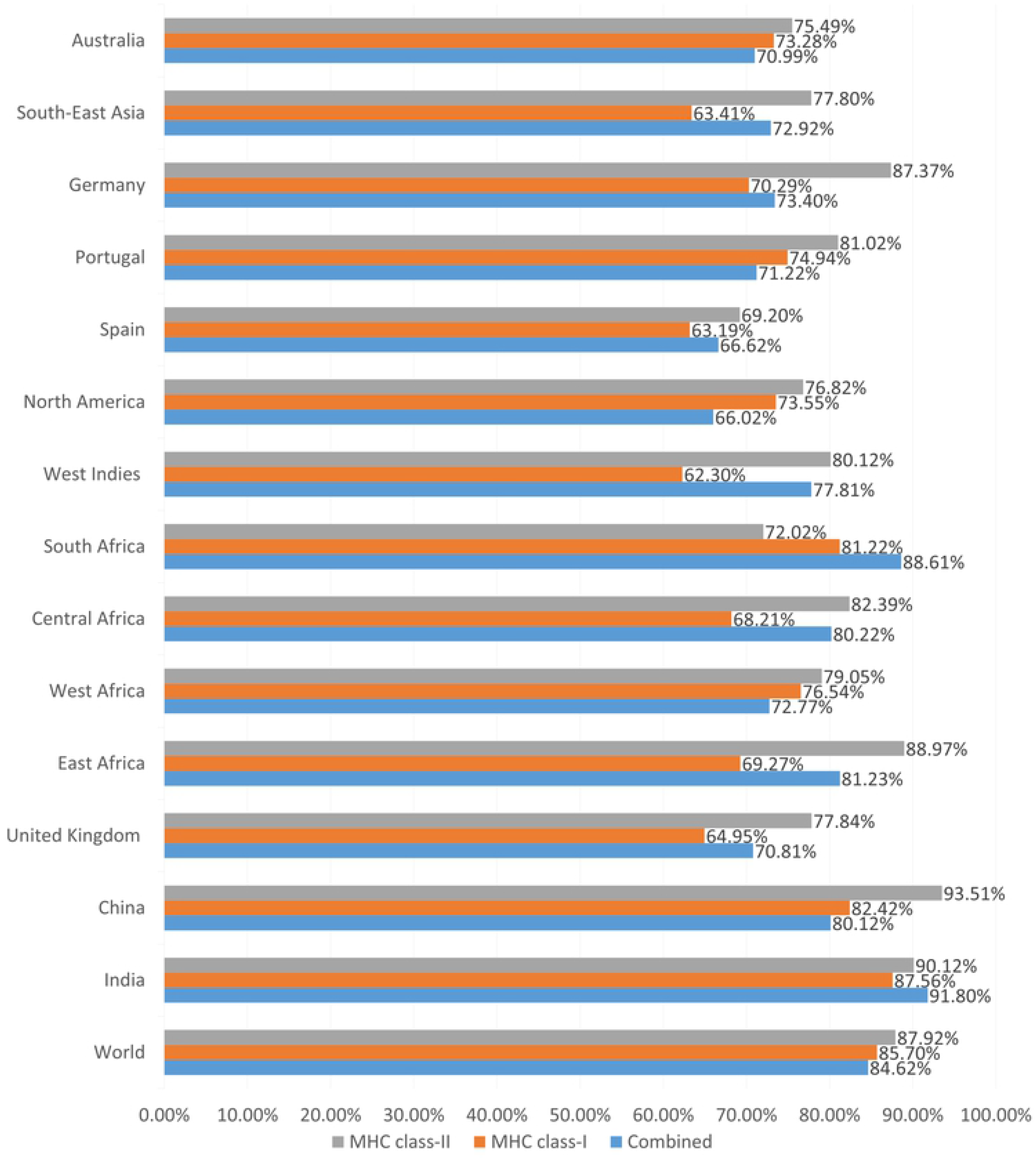
The result of the population coverage analysis of the most promising epitopes and their selected MHC alleles

Cluster analysis of the potential alleles of MHC class I and MHC class II that may interfere with the predicted epitopes of the RSV query proteins was also conducted. The study was carried out using the online tool MHCcluster 2.0, which phylogenetically demonstrates the relation of the allele clusters. **S1 Fig** shows the outcome of the experiment where a strong interaction is shown in the red zone and a weaker interaction in the yellow zone.

### 3.5. Designing of the multi-epitope subunit vaccine

The most promising T-cell and B-cell epitopes were used to design the multi-epitope vaccine incorporating adjuvant and appropriate linkers. The hBD-3 was used as an adjuvant to design the vaccine and the PADRE sequence was also used as a potent inducer of immunity. The adjuvant was linked with the epitopes by the EAAAK linker. Furthermore, AAY, GPGPG, and KK were used to associate the epitopes with each other at their appropriate positions as given in **Fig 3**.

**Fig 3.**
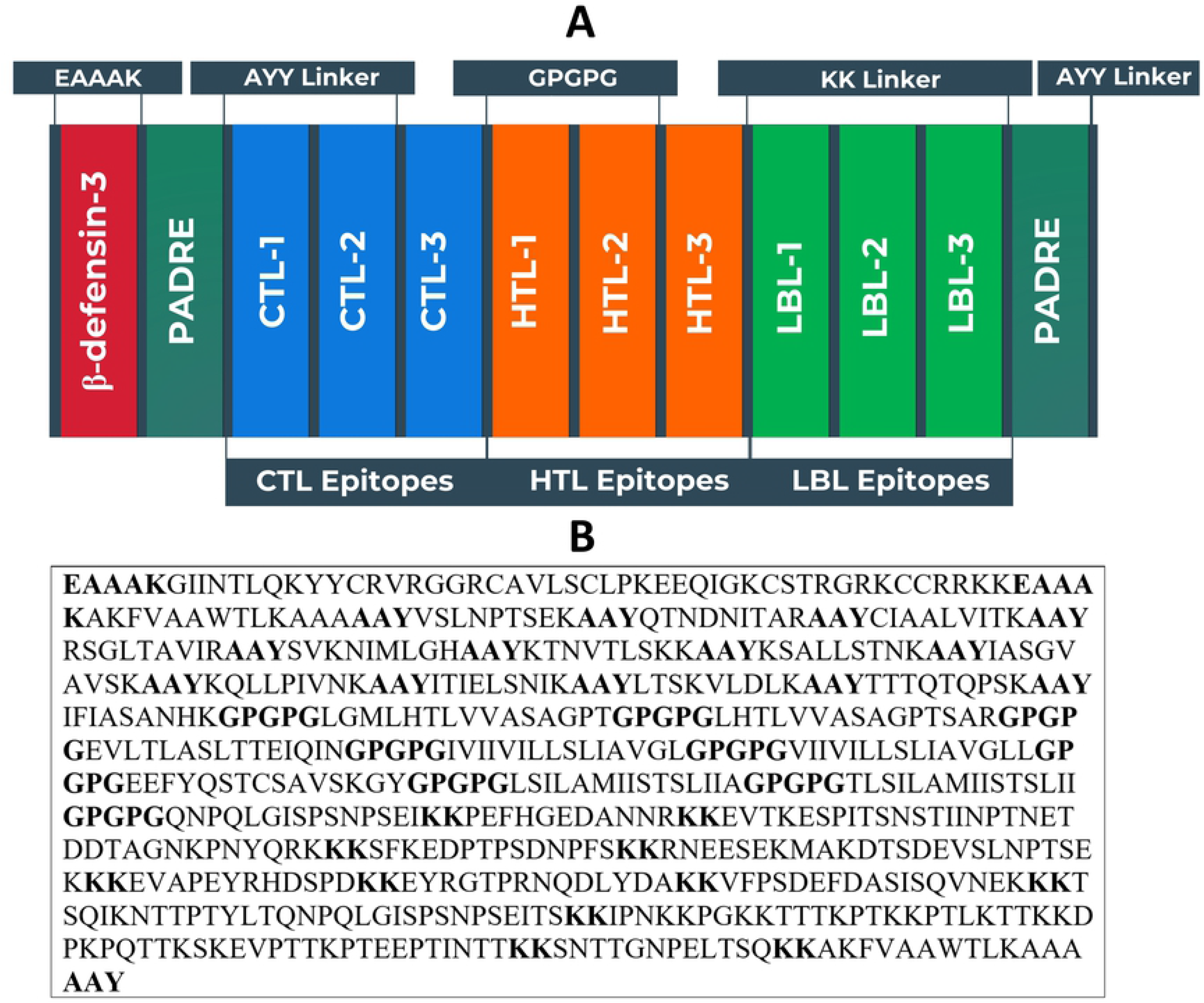
(A) Schematic representation of the potential vaccine construct with linkers (EAAAK, AAY, GPGPG, and KK), PADRE sequence, adjuvant (hBD-3) and epitopes (CTL, HTL, and LBL) in a sequential and appropriate manner (B) Sequence of the vaccine protein. The letters in bold represent the linker sequences.

### 3.6. Prediction of antigenicity, allergenicity and Physicochemical property analysis of the vaccine

The vaccine protein was observed to be both a potent antigen and a non-allergen. The vaccine had a high theoretical (basic) pI of 9.75. It had a reasonably adequate half-life in mammalian cells of 30 h and of more than 10 h in the *E. coli* cell culture system. The GRAVY value of the vaccine was considered to be significantly negative at -0.362. Additionally, both servers, Sol-Pro and Protein-sol, have also shown that the vaccine protein is soluble, attesting to its negative value. The instability index of the protein was found to be less than 40 (27.55), indicating the vaccine to be quite stable. The extinction coefficient and the aliphatic index of the vaccine were also found to be high with values, 45770 M^-1^ cm^-1^ and 80.85, respectively.

### 3.7. Secondary and tertiary structure prediction of the vaccine

The secondary structure of the vaccine protein revealed that the coil structure had the largest number of amino acids, while the β-strand showed the lowest percentage. The predictions provided by all four servers are depicted in **Fig 4**. The amino acid percentages of α-helix, β-strand, and coil structure of the vaccine protein produced from four different servers are listed in Table 03. All of the servers revealed almost similar predictions and the overall analysis also showed that the adjuvant generated potential variations in the secondary structure of the vaccine protein. The vaccine construct’s 3D structure was predicted by the RaptorX online server. The constructed vaccine protein had a surprisingly low p-value of 8.71e-05 in 4 domains, which demonstrated that the accuracy of the proposed 3D structure was significantly good. Using 1KJ6A as the template from the Protein Data Bank, the homology modeling of the vaccine construct was completed. Furthermore, the vaccine structure was modeled using Modeller, as shown in **Fig 5**, to further improve the quality.

**Fig 4.**
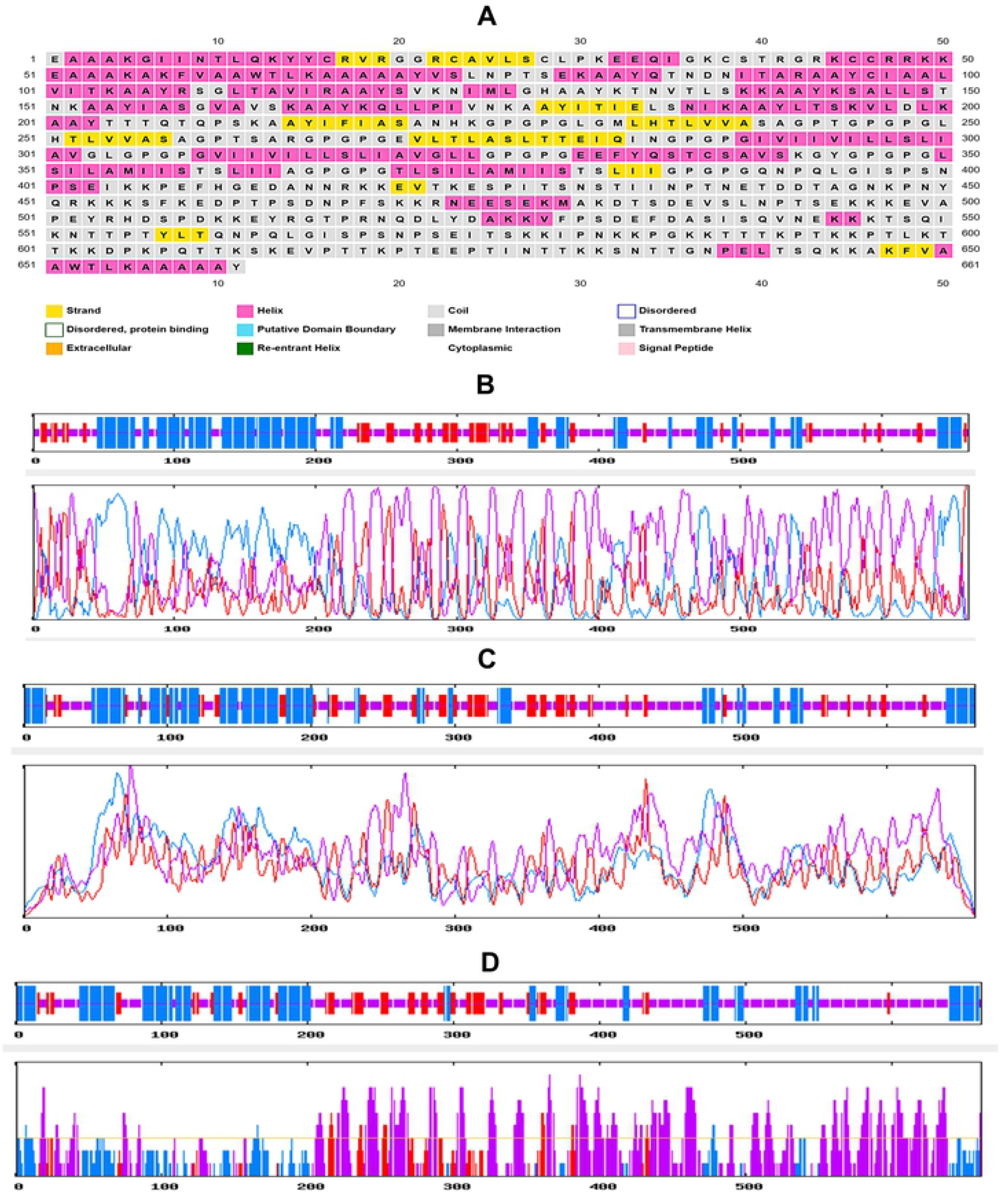
The results of the secondary structure prediction of the vaccine. (A) PRISPRED prediction, (B) GOR IV prediction, (C) SOPMA prediction, (D) SIMPA96 prediction.

**Table 03.**
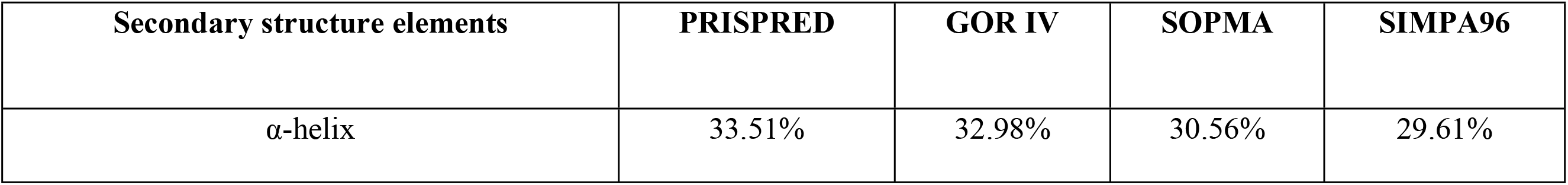
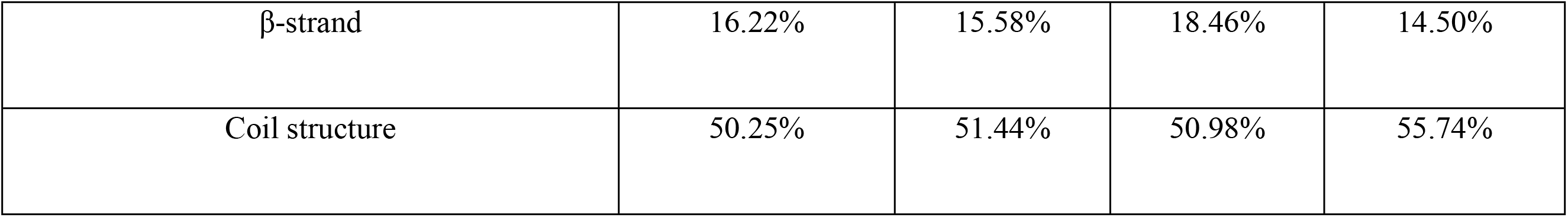
Results of the secondary structure analysis of the vaccine construct.

**Fig 5.**
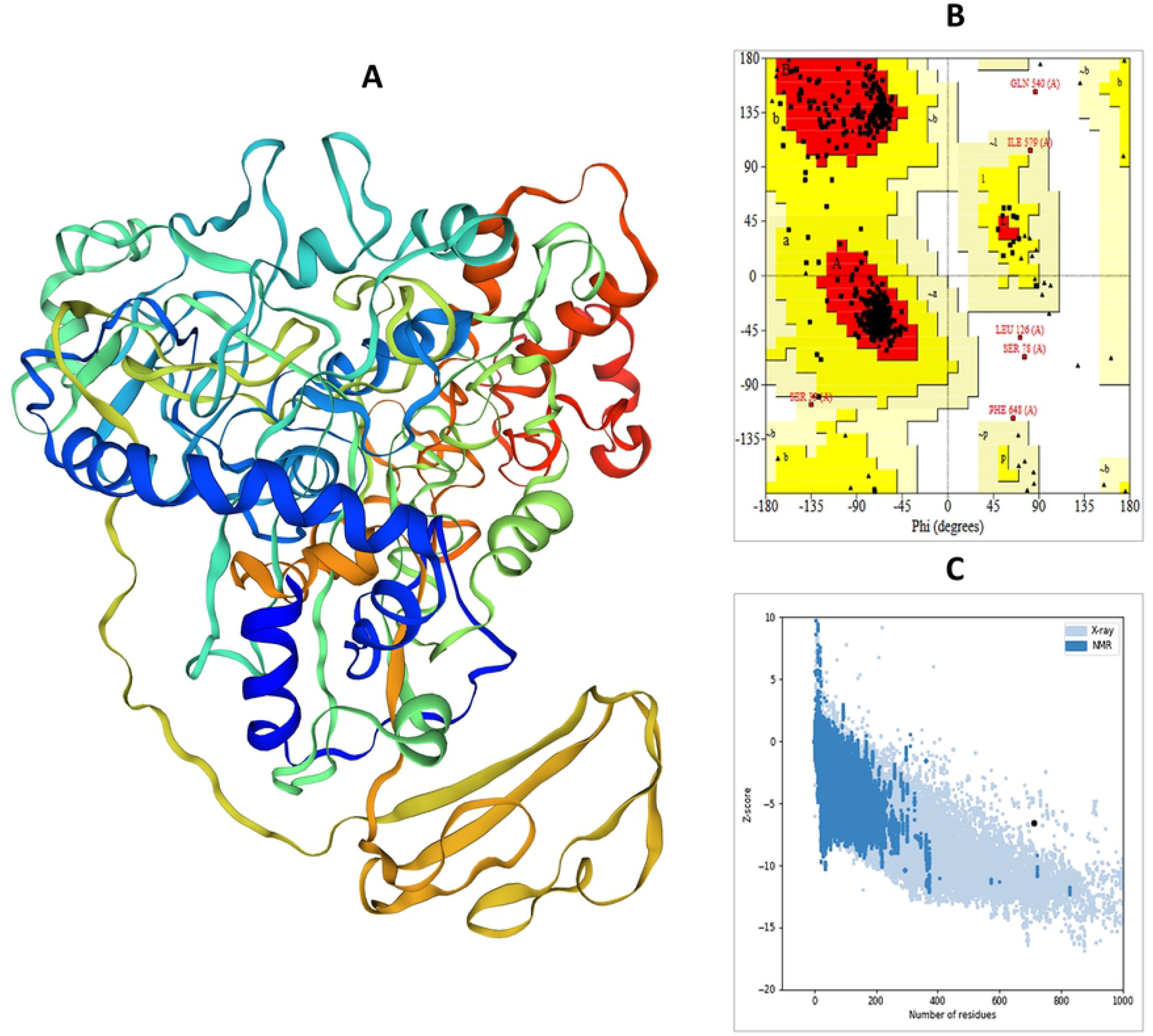
(A) The tertiary or 3D structure of the vaccine construct modeled, refined and visualized by RaptorX, GalaxyWEB server, and BIOVIA Discovery Studio Visualizer v. 17.2 respectively. (B) The results of the Ramachandran plot analysis generated by PROCHECK server and (C) quality score or z-score graph generated by the ProSA-web server of the refined vaccine construct. In the Ramachandran plots, the orange and deep yellow colored regions are the allowed regions, the light yellow regions are the generously allowed regions and the white regions are the outlier regions and the glycine residues are represented as triangles.

### 3.8. Refinement and validation of tertiary structure of the vaccine

The 3D structure of the vaccine protein produced by the RaptorX server was refined to predict a structure that closely resembles the native protein structure. The refined protein structure was then validated by evaluating the PROCHECK server-generated Ramachandran plot and the ProSA-web server-generated z-score. The Ramachandran plot study found that in the most preferred region, the vaccine protein had 93.5 % of amino acids, while in the additional approved regions, 5.4 % of amino acids, 0.4 %of amino acids in the generously permitted regions, and 0.7 % of amino acids in the disallowed regions. In comparison, the z-score of the engineered vaccine was -6.58, which is beyond the range of all experimentally confirmed X-ray crystal protein structures from the Protein Data Bank. The protein validation analysis estimated that there was a reasonably good consistency structure in the distilled form (**Fig 5**).

### 3.9. Prediction of conformational B-lymphocytic epitopes

The conformational B-cell epitopes of the vaccine protein were predicted using the ElliPro server which predicts conformational epitopes from tertiary structures of the protein. A score of 0.50 or higher was selected for the prediction of discontinuous peptides by Ellipro. Three discontinuous B-cell epitopes were predicted to include 333 amino acid residues, with values ranging from 0.506 to 0.675. The size of the conformation epitopes varied from 4 to 394 residues. Three-dimensional representation of conformational B cell epitopes of the designed multi-epitope-based RSV vaccine and the epitope residues are shown in **Fig 6** and listed in **S6 Table.**

**Fig 6.**
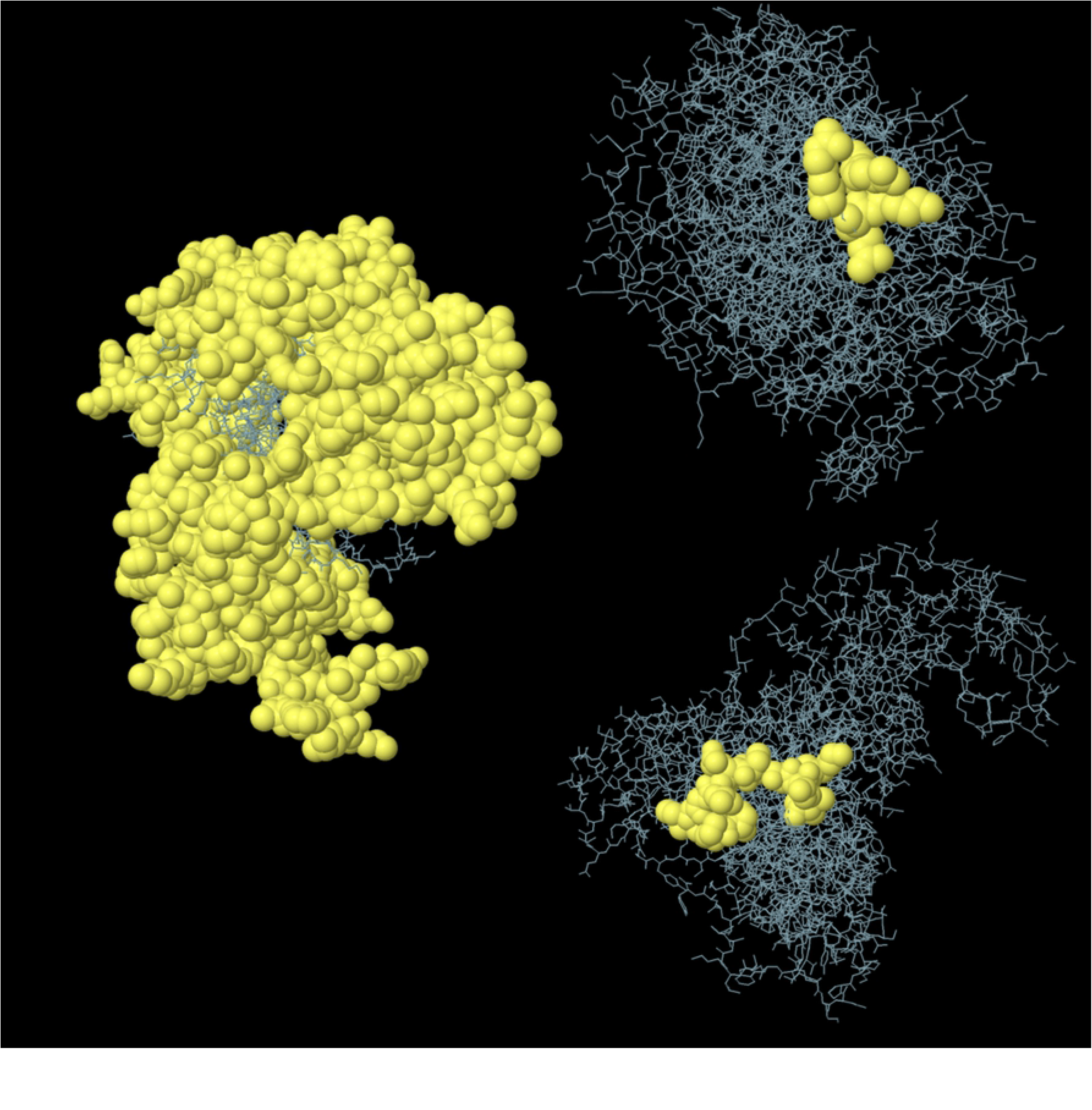
Graphical representations of the predicted conformational B-cell epitopes of the modeled vaccine indicated by yellow coloured ball-shaped structures.

### 3.10. Vaccine protein disulfide engineering analysis

The disulfide bonds of the vaccine structure were predicted using the DbD2 server in protein disulfide engineering. Based on certain classification criteria, the server recognizes pairs of amino acids with the ability to form disulfide bonds. Only those amino acid pairs that had bond energy smaller than 2.2 kcal/mol were chosen in this experiment. Three pairs of amino acids with bond energy below 2.2 kcal/mol were provided by the RSV: 23 Cys and 38 Cys, 414 Ala and 414 Lys, and 513 Tyr-516 Thr. The selected pairs of amino acids have formed the mutant vaccine in the DbD2 server, which contains potential disulfide bonds within (**S2 Fig**). This indicates the probable stability of the designed multi-epitope vaccine construct.

### 3.11. Post-translational modification analysis

The posttranslational modification analysis was performed to see whether the vaccine construct would undergo any substantial changes after being administered to mammalian cells. Four N-glycosylation sites and sixty-three O-glycosylation sites were predicted in the vaccine construct sequence. The findings suggest that a significant amount of glycosylation may have occurred inside the predicted vaccine construct, which might improve the vaccine’s efficacy and immunogenicity. In addition, the vaccine protein sequence has ninety-six phosphorylated residues (i.e., serine residues (S), threonine (T), and tyrosine (Y) phosphorylation sites) according to the NetPhos v2.0 server output. The server-provided plots containing the N-glycosylation sites and phosphorylation are given in **S3 Fig.**

### 3.12. Analysis of protein-protein docking

Protein-protein docking analysis was performed to demonstrate the vaccine’s ability to interact with various crucial molecular immune components i.e., TLRs. When docked using ClusPro 2.0, it demonstrated very high binding affinities with all its targets (TLRs). It has been further studied using the ZDOCK server where the vaccine protein also displayed very strong interaction with the TLRs. The lowest energy level obtained for docking between the vaccine construct and TLR-1, TLR-2, TLR-3, TLR-4, and TLR-9 were -986.1, -1236.7, -1084.4, -1260.8, and -1226.4, respectively. The lowest energy level between the vaccine and TLRs indicated the highest binding affinity.

### 3.13. Molecular dynamics simulation studies and MM-PBSA calculations

MD simulation is an effective method for the analysis of biological systems and it provides many mechanistic insights into the possible behavior of the system under a simulated biological environment [121]. Gromacs 2020.4 was used to carry out the production phase MD and the analysis of resulting trajectories was undertaken to understand the structural properties, and interaction between different TLRs and the predicted vaccine protein at a molecular level. The snapshots of equilibrated structures of each TLR-vaccine complex and the snapshots of the last trajectories are shown in **Fig 7**. The visual inspection of trajectories at different time intervals suggested that the side chains of the vaccine make different interactions with chain A of all TLRs except TLR1, where vaccine side chains were found to be interacting with side chains of both A and B chains.

**Fig 7.**
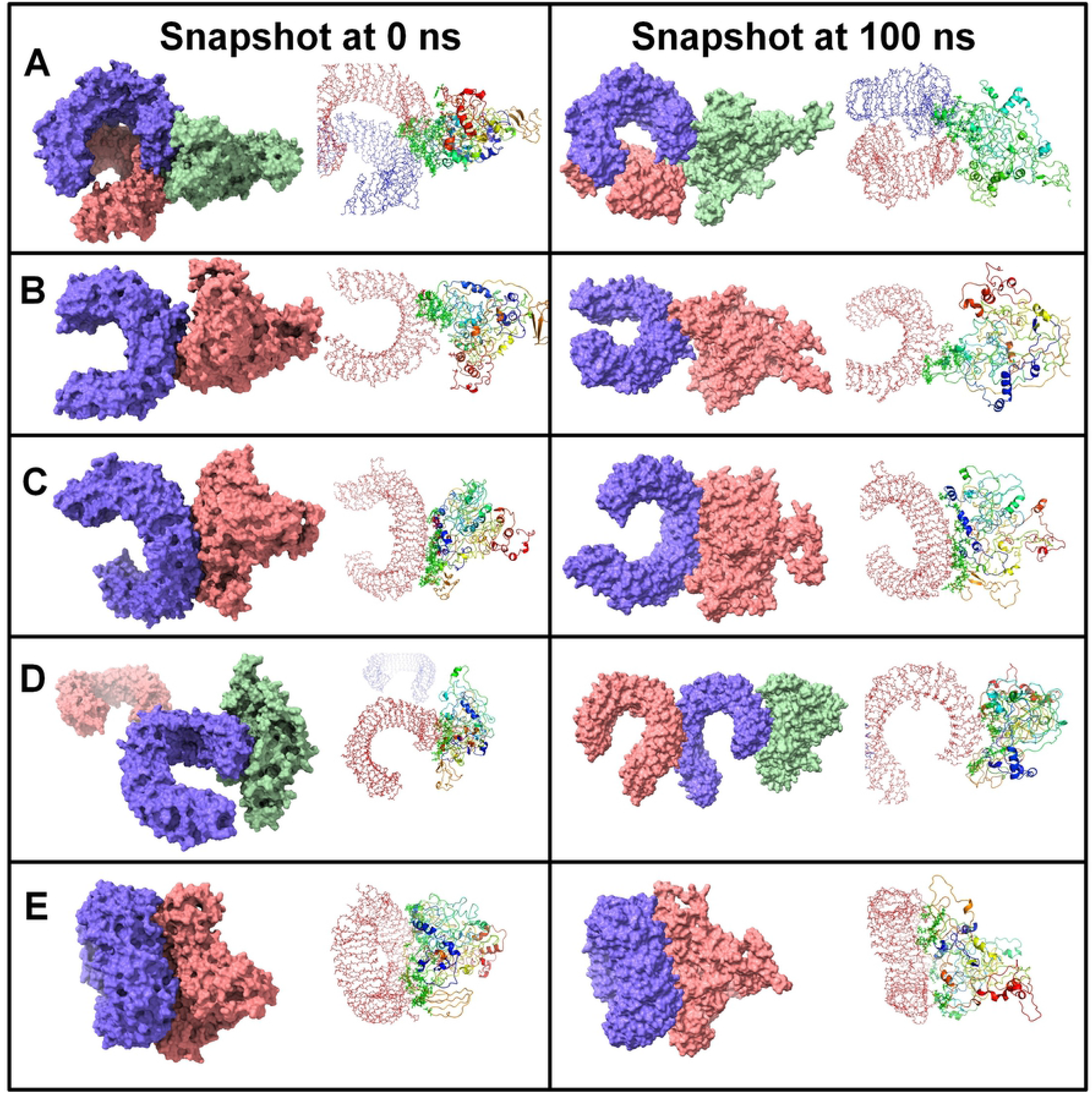
Snapshots of equilibrated (initial) systems and last trajectories. Vaccine bound complexes of A) TLR1, B) TLR2, C) TLR3, D) TLR4, and E) TLR9 (For each snapshot the surface representation and cartoon representations are shown)

Root mean square deviations (RMSD) analysis gives insights into how the backbone atoms move relative to the initial equilibrated positions. Lower the RMSD, better the stability of the corresponding system. In the present work, we measured the RMSD in backbone atoms of entire protein-protein complexes of TLRs with a vaccine. **Fig 8A** shows the RMSD in the investigated systems. The evaluation of root means square fluctuations (RMSF) provides insights into the possible changes in the secondary structure of protein under investigation. In the present work, the RMSF in the side chain atoms of residues in each system was measured. As the TLR-vaccine systems have multiple chains, the RMSF evaluation is performed on each chain of the complex to understand which residues are involved in the key contacts. The RMSF in the TLR side-chain atoms is shown in **Fig 8B**. The RMSF in other chains in each of the TLRs is given in **S4 Fig.** The analysis of radius of gyration (Rg) provides the overall measurement of compactness of the system [122]. The results of total Rg are shown in **Fig 8C**.

**Fig 8:**
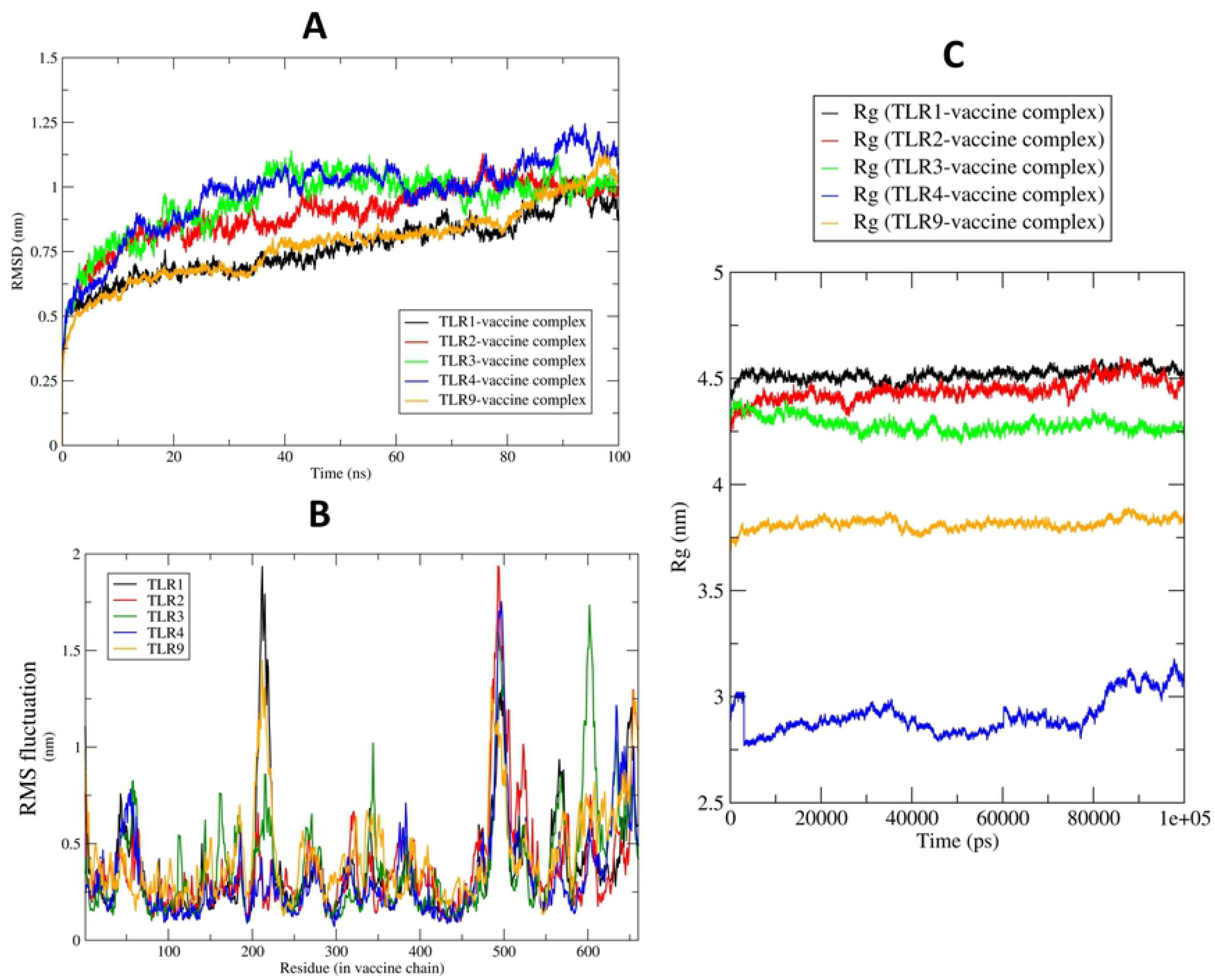
Results of the (A) Root mean square deviations in the investigated systems, (B) Root mean square fluctuations in the side chain atoms of vaccine, and (C) Radius of gyration of the vaccine.

Analysis of non-bonded interactions such as hydrogen bonds is quite challenging in protein-protein complexes. The side chains of the proteins participate in hydrogen bond interactions. The hydrogen bond analysis of all the trajectories was performed with the h-bond module of Gromacs, while the key residues at the interface of the TLR chain and vaccine chain were analyzed through the chimeraX program [123]. The results of the hydrogen bond analysis are shown in **S5 Fig**.

### 3.14. Immune simulation studies

The immune simulation study of the designed vaccine was conducted using the C-ImmSimm server which forecasts the activation of adaptive immunity as well as the immune interactions of the epitopes with their specific targets [63]. The analysis exhibited that the primary immune reaction to the vaccine could be stimulated substantially after administration of the vaccine, as demonstrated by a steady rise in the levels of different immunoglobulins i.e., (IgG1 + IgG2, and IgG + IgM antibodies) (**Fig 9A**). It was also expected that the concentrations of active B cells (**Fig 9B** and **Fig 9C**), plasma B cells (**Fig 9D**), helper T cells (**Fig 9E** and **Fig 9F**), and cytotoxic T cells (**Fig 9H** and **Fig 9I**) could steadily increase, reflecting the vaccine’s capacity to create a very high secondary immune response and healthy immune memory. However, **Fig 9G** demonstrates that the concentration of regulatory T cells would gradually decrease throughout the phases of the injections, which represents the decrease in suppression of vaccine-induced immunity by regulatory T cells [124].

**Fig 9.**
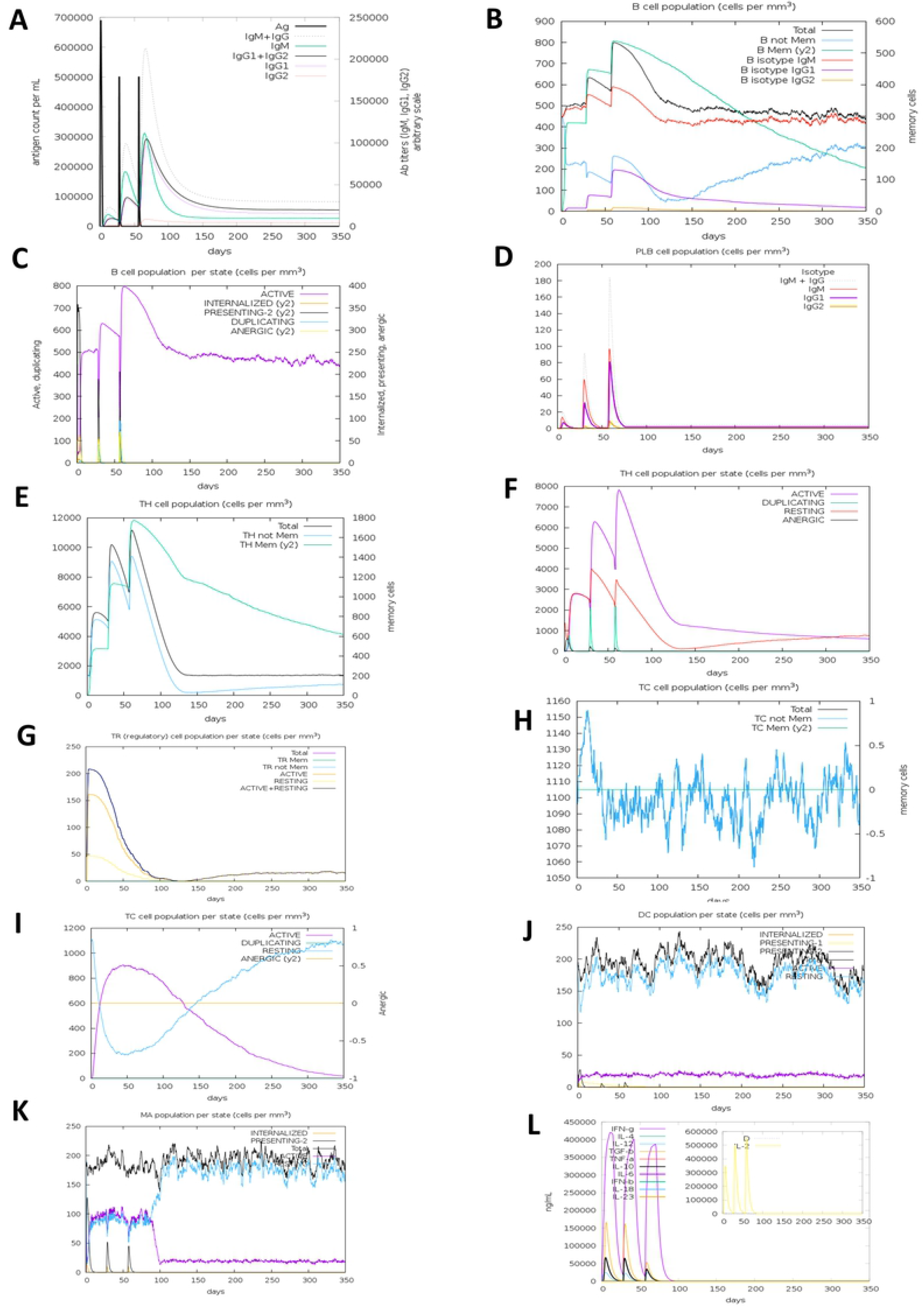

In comparison, the rise in macrophage and dendritic cell concentrations showed that these APCs had a competent presentation of antigen (**Fig 9J** and **Fig 9K**). The simulation result also predicted that the constructed vaccine could generate numerous forms of cytokines, including IFN-γ, IL-23, IL-10, and IFN-β; some of the most critical cytokines for producing an immune response to viral infections (**Fig 9L**). Therefore, the overall immune simulation analysis showed that after administration, the proposed polyvalent multi-epitope vaccine would be able to elicit a robust immunogenic response.

### 3.15. Codon adaptation, *in silico* cloning, and interpretation of the vaccine mRNA secondary structure

The protein sequence of the vaccine was adapted by the JCat server for *in-silico* cloning and plasmid construction. The CAI value was found to be 0.98, suggesting that the DNA sequences contained a larger proportion of the codons most likely to be included in the target organism’s (K12 strain of *E.coli*) cellular machinery [118, 119]. Furthermore, GC content of the formed sequence was found to be 50.23 %, within the desired range. The graph demonstrating the sequence after codon adaptation is shown in **S6 Fig**. Following codon adaptation, the projected vaccine DNA sequence was inserted between the EaeI and StyI restriction sites into the pETite vector plasmid. The plasmid includes SUMO and 6X H tag sequences that are required to promote the vaccine’s purification during downstream processing [125]. The newly built recombinant plasmid has been designated as “Cloned_ pETite” (**Fig 10**). Thereafter, the Mfold and RNAfold servers predicted the secondary structure of the vaccine mRNA. A minimum free energy score of -549.30 kcal/mol was produced by the Mfold server, which was consistent with the prediction of the RNAfold server that also predicted a minimum free energy of -526.30 kcal/mol. In **S7 Fig**, the vaccine mRNA secondary structure is depicted.

**Fig 10.**
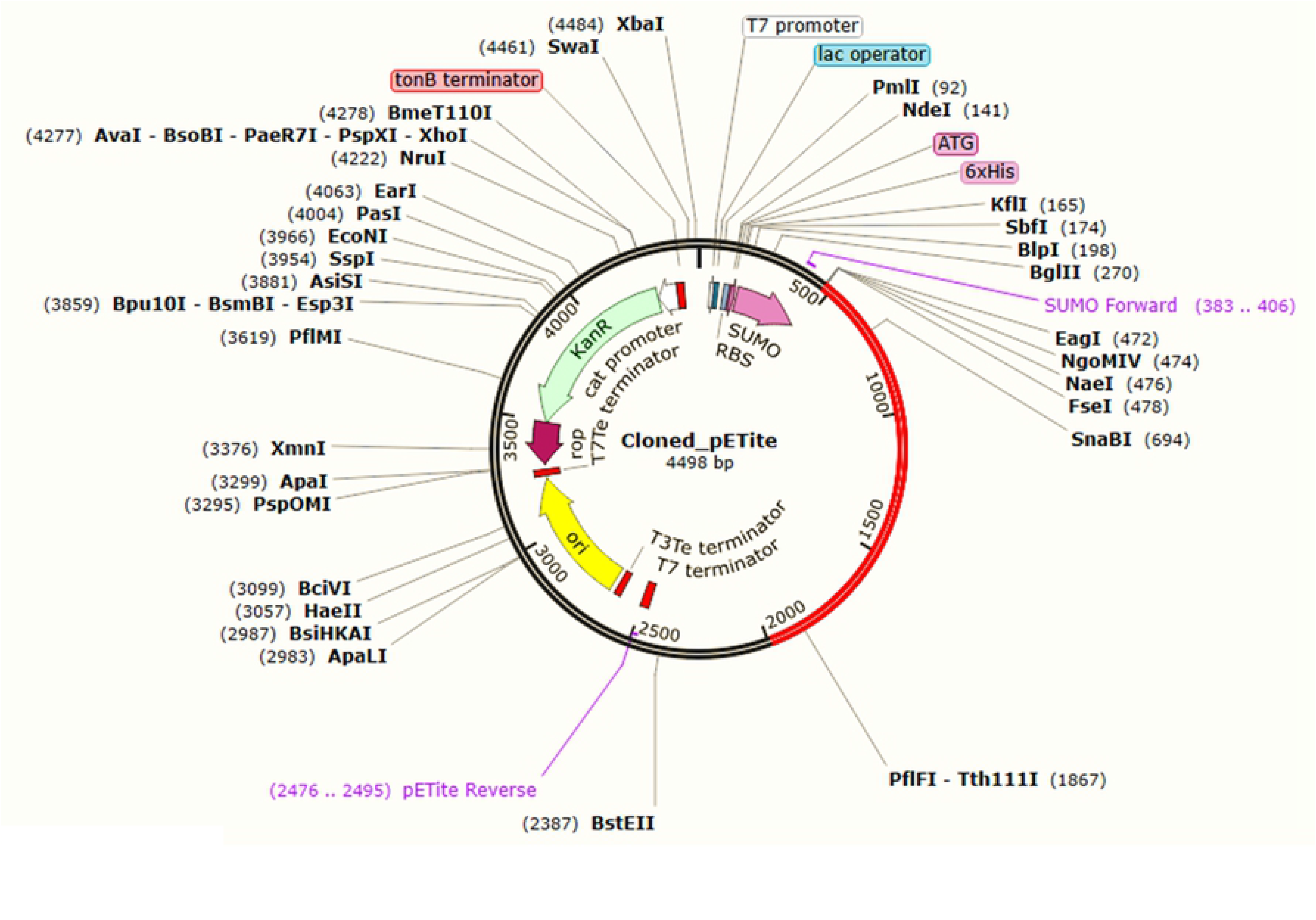
*In-silico* cloning of the vaccine sequence in the pETite plasmid vector. The codon sequence of the final vaccine is presented in red generated by the JCat server. The pETite expression vector is in black.

## 4. Discussion

hRSV is conventionally the most prevalent cause of human LRTIs, known to infect individuals from all age groups but more commonly newborns and children. Implicated infections of RSV contribute to affecting and killing numerous people all over the globe, yet no pre-existing authorized vaccine is recognized as an effective measure to prevent RSV infections. Vaccines are extensively used to control and prevent diseases caused by a variety of pathogens across the world. Conventional methods are primarily used for vaccine development and manufacture, despite the associated disadvantages of being expensive and time-consuming [126]. In contrast to traditional vaccine production strategies, today’s cutting-edge research and technology as well as the availability of knowledge about the genome and proteome of almost all viruses and organisms, facilitate the design and development of novel peptide-based subunit vaccines. Subunit vaccines have the benefit of being able to eliminate toxic and immunogenic components of an antigen during a vaccine design study, making the vaccine safe to administer in people. Subunit vaccines include a limited amount of viral particles that cause patients to develop protective immunity. A subunit vaccine is a cost-efficient and effective way to prevent health concerns [127–129]. As a result, bioinformatics and immunoinformatics techniques have been developed and widely utilized to design novel subunit vaccines that are safe, effective, efficient, and low-cost alternatives to current preventive measures [130, 131].

The narrated experiment utilized immunoinformatics methods to design a blueprint of a polyvalent epitope-based vaccine against the antigenic subgroups, RSV-A and RSV-B, targeting four distinctive proteins which include - P protein, N protein, F protein, and mG protein. Antigenicity and physicochemical properties of the proteins were predicted, where all the proteins identified were shown to be antigenic; a requirement for the use of a target protein in epitope-based vaccine construction. The theoretical pI represents the pH at which there is no net effective charge and mobility in a protein as well as predicts whether a protein is basic or acidic [132]. The instability index of a protein represents the likelihood of that specific compound being stable, and a compound with an instability index greater than 40 is deemed unstable [133]. The aliphatic index measures the relative amount of amino acids occupied by aliphatic amino acids in its side chains [134]. The high aliphatic index also indicates the improved thermal stability of a protein [135]. All of the query proteins had a high extinction coefficient and theoretical half-life of 30 h in mammalian cells. The extinction coefficient indicates the ability of a protein to absorb light at a certain wavelength and a higher extinction coefficient represents the higher absorbance of light by the protein [136, 137]. The GRAVY value represents a compound’s hydrophilic or hydrophobic traits. The GRAVY negative value reflects hydrophilic characteristics, while the GRAVY positive value reflects the hydrophobic characteristics of the compound [138, 139]. Since all query proteins were found to be quite antigenic having a high aliphatic index and extinction coefficient, as well as negative GRAVY values, they were expected to be thermostable, high light-absorbing as well as hydrophilic in nature. Overall, the physiological property analyses of the proteins revealed satisfactory results, desired for the epitope predictions. CTL, HTL, and LBL epitopes are some mandatory constituents for a multi-epitope subunit vaccine, known to stimulate or activate the cytotoxic T-cells, helper T-cells, and B-cells to generate an effective host immune response [140]. Cytotoxic T-cells can recognize the foreign antigens while helper T-cells recruit the other immune cells including B-cells, macrophages, and even cytotoxic T-cells to ensure the generation of immune responses [38, 48]. In addition, B-cells mediate the humoral immune response by producing immunoglobulins or antibodies that are antigen-specific [50][141, 142].

Once the vaccine protein reaches the host antigen-presenting cells (APC), they are processed, and the T cell epitopes are proteolytically cleaved off the protein, which is then represented by MHC molecules on the surface of APCs, exposing them to T cell receptors [143]. MHC class I molecules represent endogenous antigens often referred to as epitopes, such as intracellular proteins of a pathogen (e.g., bacteria or virus) or any tumor-inducing proteins whereas, MHC class II molecules represent exogenous epitopes. Furthermore, the antigen region that binds to the immunoglobulin or antibody is referred to as the B-cell epitope. These B-cell epitopes can be found in any exposed solvent area of the antigen and can be of various chemical types. The majority of antigens, however, are proteins, which are the targets of epitope prediction algorithms. The goal of B-cell epitope prediction is to ensure a more convenient method to identify B-cell epitopes, to substitute antigen for antibody production by the plasma B-cells, or conduct structure-function studies. Thus, antibodies can recognize any area of the antigen that has been exposed to solvents. B-cell epitopes can be split into two categories: linear and conformational; conformational B-cell epitopes are made up of patches of solvent-exposed atoms from residues that are not always sequential, while LBL epitopes are made up of sequential residues. Antibodies that identify LBL epitopes can recognize denatured antigens, but denaturing the antigen causes conformational B-cell epitopes to lose their recognition [144]. T-cell and B-cell epitopes have been predicted for the selected RSV proteins using the IEDB server. The most conserved epitopes with high antigenicity, non-allergenicity, and non-toxicity were screened for designing the vaccine construct. The broad-spectrum activity of the vaccine over the selected strains of both RSV-A and RSV-B viruses was assured by the conservancy of the epitopes.

Furthermore, the cytokine-producing ability was considered as a criterion to screen the HTL epitopes desired for designing the vaccine. Inflammatory mediators, such as cytokines and chemokines have been linked to RSV pathogenesis. They may be divided into two groups depending on how they affect immune cells: pro-inflammatory and anti-inflammatory chemicals [144]. Interleukin (IL)-1, tumor necrosis factor-alpha (TNF-α), interferon-gamma (IFN-γ), and interleukin-6 (IL-6) are pro-inflammatory cytokines [145–147]. IL-10 and IL-12 are anti-inflammatory cytokines [148, 149]. IFN-γ , on the other hand, has a dual role during RSV infection; it is essential to decrease viral multiplication while simultaneously inhibiting airway blockage [150]. IFN-γ is well-known for its fundamentally safe responses and potential to stop viral multiplication [151, 152]. Additionally, IL-4 plays important role in regulating the responses of lymphocytes, myeloid cells, and non-hematopoietic cells. In T-cells, IL-4 induces the differentiation of naïve CD4 T cells into Th2 cells, and in B cells, IL-4 drives the immunoglobulin (Ig) class switch to IgG1 and IgE, and in macrophages, IL-4, as well as IL-13, induce alternative macrophage activations [153]. Consequently, cytokines including IFN-γ, IL-10, and IL-4 were considered essential during the prediction of the HTL epitopes of the vaccine which, after administration, might play a crucial role in creating a network between immune system cells [63]. The population coverage analysis was performed in the following step. As HLA allelic distribution varies between geographical regions and ethnic groups throughout the world, it is paramount to consider population coverage while designing a viable epitope-based vaccine that is pertinent to global populations. According to the population coverage analysis, the MHC Class-I and Class-II alleles and their epitopes covered 85.70 % and 87.92 % of the global population, respectively, while the combined MHC Class-I and class-II alleles and their epitopes covered 84% of the world population. When compared to the overall population, selected epitopes exhibited a greater individual percentage cover which indicates the potential worldwide effects of the vaccine against RSV infections.

The most promising epitopes had been conjugated by specific linkers, i.e. EAAAK, AAY, KK, and GPGPG. An innate antimicrobial peptide, hBD-3 was used as an adjuvant during the construction of the vaccine. An adjuvant is considered crucial for designing a subunit vaccine because it improves the traits of antigenicity, immunogenicity, durability, and longevity of the subunit vaccine [154]. The hBD-3 was chosen as it could potentially induce TLR-dependent expression of the co-stimulatory molecules - CD80, CD86, and CD40 on the surface of monocytes and myeloid dendritic cells [155]. Moreover, by forming a protective barrier of immobilized surface proteins, hBD-3 can prevent the fusion of the virus [156]. Furthermore, it activates the APCs through TLR1 and TLR2 [157], stimulates IL-22 [158], TGF-α [159, 160], and IFN-γ [161, 162]. It also facilitates the chemotaxis of immature DCs and T cells through its interaction with chemokine receptor 6 (CCR6), as well as the chemotaxis of monocytes through its interaction with CCR2 [163]. This peptide also promotes and activates myeloid DCs and natural killer (NK) cells [157, 162].

Alongside the adjuvant, to strengthen the immunogenic reaction of the vaccine, the PADRE sequence was also incorporated. The antigenicity, allergenicity, and physicochemical properties of the constructed vaccine were subsequently identified, which revealed the vaccine protein to be desirable for further modeling refinement and validation processes. The vaccine protein was predicted with a theoretical pI of 9.75, indicating that the vaccine protein is basic and it might belong to the positively charged proteins [132]. The vaccine’s GRAVY value was predicted to be quite negative (−0.362), indicating that the vaccine protein is hydrophilic [138, 139]. Moreover, the instability index was determined to be less than 40 (27.55), implying that the vaccine is fairly stable. The extinction coefficient which is representative of the light-absorbing nature, and the aliphatic index which denotes the high stability of the vaccine protein, were both found to have high values at 45770 M-1 cm-1 and 80.85, respectively [133]. Using different online tools to predict the secondary and tertiary structure of the vaccine, it was revealed that the adjuvant sequences had produced some substantial changes in the predicted vaccine construct. Furthermore, the secondary structure analysis showed that the vaccine protein sequence was abundant with the coiled regions as well as the very low amount of β-strand. This indicates the higher stability and conservation of the predicted vaccine model. The tertiary structure was modeled and refined once the secondary structure was determined. The tertiary structure prediction of the vaccine protein revealed a p-value of 8.71e-05 in four domains of the protein, indicating that the predicted 3D structure was quite accurate. The quality of the vaccine was greatly enhanced following refinement in the context of GDT-HA, MolProbity, Rama favored amino acid percentage, and z scores, according to the tertiary structure refinement and validation study. With only a few amino acids in the outlier regions, the refined structure showed a very high Rama favored amino acid percentage. Following that, the refined structure was used for disulfide engineering. Furthermore, disulfide engineering of the vaccine construct has been conducted to increase its stability using the DbD2 v12.2 servers. The server can determine the B-factor of areas involved in disulfide bonding as well as identify potential disulfides that increase the protein’s thermal stability [92]. All residue pairings in a given protein structural model are quickly analyzed for closeness and geometry compatible with disulfide formation, assuming the residues have been changed to cysteines. Similarly, the experimental result shows residue pairings that match the specified requirements. Engineered disulfides have been shown to improve protein stability and aid in the study of protein dynamics and interactions [94]. Three pairs of amino acids have been found by the server which is predicted to improve the stability of the vaccine construct thereby. However, the vaccine’s possible effectiveness and immunological responses may be reduced if posttranslational modification, such as glycosylation and phosphorylation, is overlooked during vaccine development. Glycosylation is a chemical modification of macromolecules in which carbohydrate moieties are covalently bonded to the N or C terminals of lipids and proteins molecules, resulting in N-linked and O-linked glycosylation [164]. Previous studies have found that glycosylation significantly improves vaccine immunogenicity when compared to non-glycosylated vaccinations, and clinical trials are now ongoing [165]. Furthermore, phosphorylation is the process of adding a phosphate group to macromolecules. Eukaryotes have a greater frequency of occurrence of posttranslational modifications. Serine and threonine are two important phosphorylation sites. Phosphorylated peptides or epitopes (synthetic or natural) are known to be better recognized by cytotoxic T cells, i.e. MHC Class I molecules, and are therefore directly implicated in the production of particular immune responses [166]. As a result, it is hypothesized that the vaccine construct predicted with multiple posttranslational modifications will create an efficient immunogenic response against the virus following vaccine administration.

Furthermore, one of the predominant necessities and approaches to designing an effective vaccine is the molecular docking process. It concludes the probability of contact between the vaccine and other networking proteins, i.e. TLRs may occur during initial immune response. TLRs, which are found on leukocytes and in tissues, play a major role in innate immunity activation by identifying invading pathogens, including viruses like RSV, and sending out signals that promote inflammation-related components [167]. TLRs, such as TLR2, TLR1, TLR6, TLR3, and TLR4, are found on leukocytes and can interact with RSV to boost immune responses [168]. Within the lungs, TLR2 interactions with RSV increase neutrophil migration and dendritic cell activation. TLR2 exists as a heterodimer complex with either TLR1 or TLR6 on the surface of immune cells and tissues [169]. According to genetic analysis and vaccine studies, TLR2 signaling appears to be critical in RSV recognition [170–172]. TLR2 and TLR1 or TLR2 and TLR6 complexes can recognize RSV, and greatly enhance early innate inflammatory responses [173–175]. Previous research has also suggested that the signals generated by TLR2 and TLR6 activation are critical for viral replication control [168]. TLR3 and TLR4 signaling support the T helper type 1 (Th1) responses, whereas T helper type 2 (Th2) responses are favored by TLR2/1, and TLR2/6 signaling [177]. Th1 cells help with responses to intracellular pathogens, whereas Th2 cells handle parasitic infections and allergies [178, 179]. The F glycoprotein of RSV has been reported to induce primarily a Th1-type immune response through the interaction with TLR4 [180, 181]. On the other hand, TLR9 has been found to improve vaccine immunogenicity and decrease vaccine-enhanced illness during FI-RSV immunization. Furthermore, immunomodulation generated by TLR9 agonists verifies TLR agonists’ adjuvant potential after RSV immunization [182]. Finally, the targeted TLRs are all involved in pattern recognition and the innate immune response to RSV, which results in the production of proinflammatory cytokines and chemokines [183]. TLRs which are critical in the RSV pathogenesis were considered for the docking analysis. Thus, TLR-1, TLR-2, TLR-3, TLR-4, and TLR9 have been docked with the vaccine protein which revealed each of the TLRs to obtain very low binding energy in different servers and to show strong interaction with the vaccine protein according to the results given in the docking analysis. It is evident from the docking analysis that the designed vaccine would have a strong affinity with all the target TLRs, leading to the possibility that a strong immune response might be induced by the vaccine after administration.

In response to external forces exerted by its surrounding environment, MD simulation examines the motion and changes in the state of a target protein molecule or complex. In this experiment, to get a better insight into their molecular stability, the five docked vaccine-TLR complexes were simulated. The experiment revealed that the structures retained appropriate levels of deviation. The TLR1-vaccine complex was found to have the least RMSD compared to other complexes; the TLR4-vaccine complex had higher RMSD despite having a similar number of chains. The result of RMSD represents the stability of TLR1 and TLR9 vaccine complexes. Despite 3 chains *viz.* Chain A, chain B, and vaccine chain in the TLR1-vaccine complex, the resultant RMSD, suggest reasonable stability. Furthermore, The RMSF analysis suggests the residues in the range 190-240 and 480-530 are having large magnitudes of fluctuations in all the complexes. The TLR3-vaccine complex showed slightly larger fluctuating side chains than other complexes. The results of the total Rg analysis suggest that the total Rg of the TLR1-vaccine complex almost remained constant throughout the simulation. The total Rg of the TLR4-vaccine complex was found lower; however, it deviated throughout the simulation. Though both these TLRs have two chains of proteins along with vaccine chains, TLR1 Rg seemed quite stable, while the other TLR systems (TLR2, TLR3) also seemed to be quite compact, and there were no evident secondary structural changes in these TLRs. In the non-bonded interaction analysis, in the case of TLR1-vaccine complex, it is found that around 10 hydrogen bonds were formed till around 50 ns MD time interval, which steadily lowers to around 5 hydrogen bonds till 80 ns and thereafter rises to 10 hydrogen bonds. The TLR2-vaccine complex has around 7 hydrogen bonds being constantly formed throughout the MD simulation, while TLR3 and TLR9 complexes with vaccines have around 10 hydrogen bonds constantly formed. These complexes have strong hydrogen bond networks at the interface of TLRs and vaccines. The TLR4-vaccine complex has around 5 hydrogen bonds formed which are fewer in number compared to other systems. Actual residues involved in the hydrogen bond formation as investigated in the last trajectory are tabulated in the **S7 Table.**

The immune simulation analysis of the proposed vaccine demonstrated that the vaccine could induce an immune response compatible with the natural host immune system. Both humoral and cell-mediated responses may be activated by the vaccine, as shown by an elevation in the levels of memory B cells, plasma B cells, cytotoxic T cells, helper T cells, and various antibodies. Adaptive immunity is an immunity that occurs after exposure to an antigen either from a pathogen or a vaccination. A vaccine can generate adaptive immunity against the pathogen by which it may restrict or prevent the infection. The vaccine-provided activation of helper T cells resulted in strong adaptive immunity [184–186]. Again, a very strong antigen presentation was also demonstrated in the simulation study by the rise in the concentration of APCs such as macrophages and dendritic cells. In addition, enrichment in the cytokine profile that plays a crucial role in providing broad-spectrum immunity against viral invasions [186–189] has also been identified in the analysis [66]. Moreover, the gradual increase in the level of different mucosal immunoglobulins i.e. IgG1 + IgG2, and IgG + IgM antibodies throughout the vaccine doses or injections were also predicted. The mucosal immune system is the dominant part of the immune system, having developed to protect the mucosae in the upper respiratory tract, which are the primary sites of respiratory infection. Being a respiratory virus, RSV initially affects the upper respiratory tract, due to which the immune system may be stimulated against the virus predominantly at the mucosal surfaces [190]. Previous clinical studies also reported IgG and IgM to be significant role players against RSV infections [191]. Furthermore, the simulation analysis revealed that the concentration of regulatory T cells would gradually decrease throughout the phases of the vaccine doses which indicates the potential decrease in suppression of vaccine-induced immunity by regulatory T cells [192].

Hence, the proposed vaccine construct is predicted to produce an effective immunogenic response after the vaccine injections, according to the analysis of the immune simulation. The codon adaptation and subsequent *in silico* cloning studies were conducted to identify the potential codons required for the generation of a recombinant plasmid that could be used to express the vaccine in the *E. coli* strain K12, leading to the mass manufacturing efforts of the vaccine in the near future. The *E.coli* cell culture system is considered to be the majorly recommended system for the production of recombinant proteins at a mass level. In the codon adaptation analysis, the obtained results were significantly good with a CAI value of 0.98 and a GC content of 50.23 %, since any CAI value above 0.80 and a GC content of 30% to 70% are considered to be the most promising scores [34, 186, 193]. Following this, the optimized vaccine DNA sequence was inserted into the pETite plasmid vector using Snapgene restriction cloning software. The pETite plasmid vector contains SUMO tag and 6X His tag which might be fused with the vaccine codon sequence during the *in silico* cloning process. This may lead to the expression of these tags within the protein itself, which could promote the purification and downstream processing of the vaccine. Prediction of the stability of the vaccine mRNA secondary structure using the Mfold and RNAfold servers provided with the negative and much lower minimal free energies of -549.30 and -526.30 kcal/mol, respectively. The lower minimal free energy is often considered better than the higher maximal free energy score which indicates the protein to be more stable. It can, therefore, be reported that the predicted vaccine could be very stable upon transcription [133]. Overall, this study suggests that the proposed vaccine peptide could be utilized as a potential and successful protective measure against both RSV-A and RSV-B subtypes. However, to eventually validate its immunogenicity, efficacy, stability, safety, and various biophysical characteristics, further research approaches, and implementations are recommended.

## 5. Conclusion

RSV is predominantly one of the major contributors of diseases of the LRTI, including pneumonia and bronchiolitis, responsible for infecting people of all ages as well as immunocompromised individuals with a high infection rate. Millions of individuals eventually end up being diagnosed with this virus every year, and a huge proportion of them require to be hospitalized. While research for an effective countermeasure to tackle this virus has been ongoing for the past few decades, no approved vaccine is still commercially available. Moreover, the antiviral drugs now available often struggle to show any effective outcomes during therapy. Therefore, a potential epitope-based polyvalent vaccine against both forms of RSV, RSV-A, and RSV-B, was designed in this research using the techniques of immunoinformatics and *in-silico* biology. The vaccine included the T-cell and B-cell epitopes that were 100% conserved; they could therefore be efficient against the two selected viruses. In addition, high antigenicity, non-allergenicity, and non-toxicity as well as non-homology (to the human proteome) were also considered to be the criteria for choosing the most promising epitopes for the final construction of the vaccine, so that the vaccine could deliver a very strong immunogenic response without triggering any adverse reaction inside the body. The results of the various analyses conducted in the study revealed that the polyvalent vaccine should be very safe, efficient, and responsive to use. The tools utilized in this study are well accepted and yield highly accurate results. Therefore, the findings of this study can point researchers in the direction of novel vaccine development tactics. Researchers could investigate the predicted epitopes and their probable immunogenic response elicited in the host system when looking into further subunit vaccine development or other prevention strategies against RSV infection. However, as all these predictions were focused solely on computational techniques, it is important to perform further wet laboratory-based experiments to validate the findings of this analysis. With high-cost criteria and numerous drawbacks in improving the preparation of a live, attenuated, or inactivated vaccine for such highly infectious agents, candidates for peptide-based vaccines, such as the one designed in this study, maybe comparatively cheap and an efficient alternative to reach the entire world as a polyvalent vaccine to fight the challenge of RSV infections.

## 6. Acknowledgements

Authors are thankful to the Department of Pharmaceutical Chemistry, Sinhgad College of Pharmacy, Maharashtra, India, Department of Genetic Engineering and Biotechnology, University of Chittagong, Chattogram, Bangladesh and Swift Integrity Computational Lab, Dhaka, Bangladesh, a virtual platform of young researchers, for providing support to successfully carry out the research.

## 7. Declarations

### Ethics approval and consent to participate

Not Applicable

### Consent for publication

Not Applicable

### Availability of data and material

All the data generated during the experiment are provided in the manuscript/supplementary material.

